# Dysfunctional β-cell longevity in diabetes relies on energy conservation and positive epistasis

**DOI:** 10.1101/2024.03.18.585508

**Authors:** Kavit Raval, Neema Jamshidi, Berfin Seyran, Lukasz Salwinski, Raju Pillai, Lixin Yang, Feiyang Ma, Matteo Pellegrini, Juliana Shin, Xia Yang, Slavica Tudzarova

## Abstract

Long-lived PFKFB3 expressing β-cells are dysfunctional cells because of prevailing glycolysis that compromises metabolic coupling of insulin secretion. Their accumulation in type-2 diabetes (T2D) appears to be related to the loss of apoptotic competency of cell fitness competition (CFC) that maintains tissue function by favoring constant selection of healthy “winner” cells. To investigate how PFKFB3 can disguise the competitive traits of dysfunctional “loser” β-cells, we analyzed the overlap between human β-cells with bona-fide “loser signature” across diabetes pathologies utilizing the HPAP scRNA-seq and spatial transcriptomics of PFKFB3 positive β-cells from nPOD T2D pancreata. The overlapping transcriptional profile of “loser” β-cells was represented by downregulated ribosomal biogenesis- and genes encoding for mitochondrial respiration. PFKFB3 positive “loser” β-cells had reduced expression of HLA Class I and II genes. Gene-gene interaction analysis revealed that PFKFB3 *rs1983890* can interact with anti-apoptotic gene *MAIP1* implicating positive epistasis as a mechanism for prolonged survival of “loser” β-cells in T2D. Inhibition of PFKFB3 resulted in the clearance of dysfunctional “loser” β-cells leading to restored glucose tolerance in mouse model of T2D.

## INTRODUCTION

Despite the clear link between accumulation of injured and dysfunctional β-cells and type-2 diabetes (T2D), it is difficult to target specifically these pathogenic cells in a therapeutic attempt to modify the trajectory of T2D. Recent studies proposed that long-lived dysfunctional β-cells confine to a senescent phenotype implicating them in the underlying pathology of both type-1 diabetes (T1D) and T2D (Aguayo-Mazzucato *et al*, 2019; Moiseeva *et al*, 2022; Thompson *et al*, 2019). However, targeting senescent cells is a daunting task because of their notorious heterogeneity (Kirschner *et al*, 2020; Kwiatkowska *et al*, 2023) and the current inability to distinguish their biological benefit from their pathologic effect.

Rather than relying on the idea of pathogenic cells’ identity, we reasoned that a more effective approach would be to probe the existence of a context-dependent tissue property in health to which we can attribute the specific recognition and clearance of the injured and/or dysfunctional cells. As such, cell fitness competition (CFC), is emerging as a tissue clearance mechanism that can maintain function by monitoring tissue behavior at the population scale (Merino *et al*, 2016) during and after development (de la Cova *et al*, 2004; Gibson & Perrimon, 2005; Moreno *et al*, 2002) and in adult post-mitotic tissues (Coelho & Moreno, 2019; Coelho *et al*, 2018; Vieira *et al*, 2023). In essence, it is a homeostatic process that embodies the constant replacement of aged, injured, or dysfunctional (“loser”; CFC terminology) cells with healthier, functional (“winner”) cells (reviewed in (Bowling *et al*, 2019)). The process relies on the differential fitness within apparently isogenic cell population across several levels of cellular organization—from cellular resources (ribosomal biosynthesis, RiBi) to energy (metabolism) and infrastructure (mitochondria) (Coelho & Moreno, 2019; Coelho *et al*., 2018). Under homeostatic conditions, CFC enriches the tissue with functional and healthy cells without marked changes in tissue mass which is deemed a silent phenotype (Blaauw *et al*, 2010; Leychenko *et al*, 2011; Sasai *et al*, 2010; Tamori & Deng, 2014). When damaged beyond repair, a cell’s “molecular fitness fingerprint” is marked by protein aggregates and oxidative stress (Baumgartner *et al*, 2021). A recent body of evidence has revealed that mitochondrial dysfunction is common to different “loser” cells and that it is sufficient and necessary to trigger CFC during early mouse development (Lima *et al*, 2021). As such, “loser” epiblast cells undergo a transcriptional program in response to impaired mitochondrial function that involves integrated stress-(ISR) and unfolded protein response (UPR), implicating DNA Damage Induced Transcript 3 (*Ddit3*), Activating Transcription Factor 3 (*Atf3*), Protein Phosphatase 1 Regulatory Subunit 15A (*Ppp1r15a*) (Lima *et al*., 2021) and NFE2 Like BZIP Transcription Factor 2 (*Nfe2l2*) to restore cellular homeostasis (Melber & Haynes, 2018; Munch, 2018; Rosario *et al*, 2020). By breaking down the differentially expressed genes of the “loser-to-winner” trajectory, it was revealed that the targets of RICTOR, MYC, MYCN, and IGFR and the genes related to ribosomal synthesis primarily fell within the down-regulated genes, constituting the “loser” fingerprints (Lima *et al*., 2021).

Our previous work has demonstrated that β-cells from T2D donors have a wide range of abnormalities that mirror those that are found in molecular “loser fingerprints”, such as the proteotoxicity after accumulation of islet amyloidogenic pancreatic polypeptide (IAPP) aggregates and mitochondrial attenuation (Montemurro *et al*, 2019). Injured β-cells closely resemble “loser” cells that, however, by increasing aerobic glycolysis through the activation of the 6-phosphofructo-2-kinase/fructose-2,6-bisphosphatase 3 (PFKFB3) (Min *et al*, 2022; Montemurro *et al*., 2019; Nomoto *et al*, 2020), can escape removal by CFC (Baker, 2020; Bowling *et al*., 2019; Lawlor *et al*, 2020). Strikingly similar, pyramidal neurons in the hippocampus from sporadic Alzheimer’s disease (AD) and AD-directly converted neurons phenocopy β-cells in T2D, switching to overt glycolytic metabolism in response to Aβ proteotoxicity and oxidative stress (Traxler *et al*, 2022). This sequence of observations implies that proteotoxicity, CFC, and functional cell regeneration could be linked, with T2D and neurodegeneration (e.g., AD) potentially representing diseases of failed clearance of injured cells.

Our previous work pointed to the potential link between PFKFB3 and CFC in the human-like murine model of T2D following β-cell depletion of PFKFB3 [*PFKFB3^βKO^* on *diabetogenic stress, DS, proteotoxic stress (;hTG) and high-fat diet]*. We reported that PFKFB3 knockout led to an initial β-cell apoptotic competency and almost complete clearance of injured β-cells followed by near-wild-type levels of β-cell replication. The glucose tolerance restoring phenotype of the *PFKFB3^βKO^* DS mouse (under moderate DS stress) in connection to the elimination of injured (“loser”) cells closely reflected CFC reactivation, implicating PFKFB3 as a gatekeeper of CFC (Min *et al*., 2022).

We used dual criteria to investigate human “loser” β-cells based on 1) a *bona fide* “loser” signature adopted from the mouse embryo (Lima *et al*., 2021) and 2) PFKFB3 expression from our spatial transcriptomics analysis. We found no evidence for differential stress response between “loser” and “winner” β-cells. A notable exception was the general downregulation of genes coding for ribosomal proteins and mitochondrial respiration, which were the most common “loser signals” that eclipsed the impact of any other insufficiencies. Thus, PFKFB3 positive β-cells demonstrated a complete transcriptional overlap with “loser” ;-cells following extreme energy conservation. Given that PFKFB3 expressing “loser” β-cells are long-lived cells in T2D, we used SNP-SNP interaction analysis of PFKFB3 polymorphism *rs1983890* to unravel a genetic basis of the aborted β-cell competition in diabetes. We found that *rs1983890* interacts with collective SNPs from ArfGAP With GTPase Domain, Ankyrin Repeat And PH Domain 1 (*AGAP1*) and matrix (m)-AAA peptidase interacting protein 1 (*MAIP1*). *MAIP1* is an anti-apoptotic gene from the mitochondrial matrix that prevents constitutive activation of mitochondrial Ca^2+^ uniporter (MCU) (Opalinska & Janska, 2018).

We propose that the ribosomal biosynthesis and mitochondrial respiration represent a ‘global’ phenotypic interface of β-cell fitness that in the presence of PFKFB3 can create an epiphenomenon of positive epistasis. This epistatic regulation along with the reduced HLA expression promotes survival of dysfunctional β-cells (MacLean *et al*, 2010; Perfeito *et al*, 2014; Schoustra *et al*, 2016). We demonstrated clearance of dysfunctional β-cells and restored glucose tolerance in the murine model of T2D after inhibition of PFKFB3. These results demonstrate the feasibility of restoring islet function in T2D by unlocking CFC with small molecule PFKFB3 inhibition.

## RESULTS

### Elucidating the “loser” β-cell trajectory in diabetes

Selective elimination of PFKFB3 positive β-cells by reactivation of CFC holds potential for islet enrichment by functional β-cell regeneration **(Fig 1A)**. Therefore, we investigated a conceptual paradigm that diabetogenic stress counters the clearance of dysfunctional β-cells by a PFKFB3-dependent mechanism (**Fig 1A**). β-cell dysfunction is linked to proteotoxicity and oxidative stress, interestingly, phenocopying the “loser” cell status established previously (Baumgartner *et al*., 2021). We undertook a comprehensive genomic analysis of two independent datasets (67 donors from HPAP and 3 donors from nPOD) (**Fig 1B**). Datasets of differentially expressed genes (DEGs) were obtained by module score-integrated R script based on *bona fide* “loser” genes: *DDIT3, ATF3, PPP1R15A, RICTOR,* and *NFE2L2* (Lima *et al*., 2021) across the disease spectrum from control (Control^HPAP^), prediabetes (AAB^HPAP^), to Type-1 diabetes (T1D^HPAP^), and to Type-2 diabetes (T2D^HPAP^) and including PFKFB3 expressing ;-cells (T2D^PFKFB3^) (genomic pipeline, **Fig 1B**). Interestingly, this combined approach yielded hundreds to thousands of DEGs in all disease states except T1D, where DEGs (adjusted p<0.05) were represented by a skewed list and were not submitted to GSEA. The T1D^HPAP^ DEG list comprised genes involved in immunity such as *CD81*, immunoproteasome component *PSMB8*, cell structure regulators: mitotic spindle organizing protein (*MZT2B*), a part of gamma-tubulin complex; mitochondrial translocase *TOMM6*, microsomal glutathione-S-transferase 3 (*MGST3*), the receptor for endocytosis (*RNASEK*), TATA-box binding protein *TAF10*, tetratricopeptide repeat protein 32 (*TTC32*) and others (**Table S3**). In addition, Control^HPAP^ didn’t yield significant enrichment in any pathway (**Table S1-S5**). For the geospatial profiling, we have applied segmentation to document sequenced genes from 36-48 groups, each comprised of 100 regions of interest (ROIs) of PFKFB3 positive- and PFKFB3 negative ;-cells (**Fig EV1A-F**). Principal component analysis (PCA) predictably indicated proximity between the two β-cell subpopulations discriminated only by PFKFB3 expression (**Fig EV1G**).

**Figure 1.**
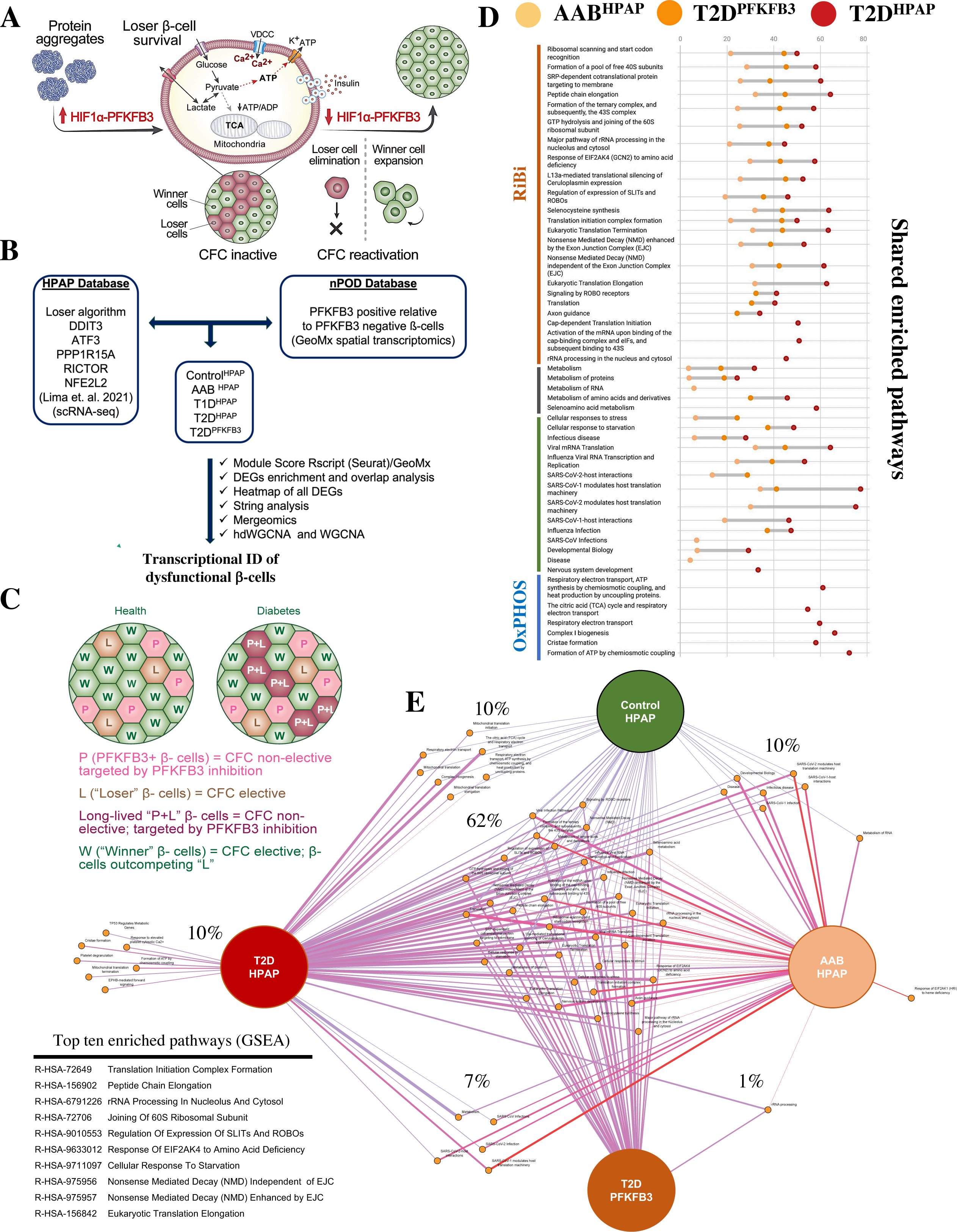
Elucidating the “loser” ;-cell trajectory in diabetes. A Scheme of PFKFB3 dependent loss of ;-cell function and locked CFC impeding functional ;-cell regeneration. Protein aggregates lead to upregulation of the HIF1α-PFKFB3 pathway that delinks glucose sensing from mitochondrial respiration and insulin secretion. PFKFB3 expression promotes “loser” ;-cells survival by CFC deactivation. Targeting PFKFB3 re-activates CFC and clearance of “loser” ;-cells. B Genomic pipeline to reveal transcriptional profile of “loser” β-cells and PFKFB3 expressing dysfunctional human β-cells across health, prediabetes (AAB), T1D, and T2D utilizing HPAP scRNA-seq database and the spatial transcriptomics from nPOD T2D pancreata, respectively. C Diagrammatic presentation of the anticipated effect of PFKFB3 targeting of “loser” ;-cells in healthy versus diabetic islets. “Loser” (L) and “Winner” (W) ;-cells are elective for CFC and CFC is active in health. However, in diabetes, “Loser” ;-cells that express high PFKFB3 (P+L) are no more CFC elective. When PFKFB3 is targeted, CFC becomes specifically unlocked in P+L cells, while ;-cells expressing low PFKFB3 (P) are not affected in the absence of L signatures. D Dot plot showing the coverage of Reactome enriched pathways of DEGs for AAB^HPAP^, T2D^HPAP,^ and T2D^PFKFB3^. E Overlap of enriched pathways across Control^HPAP^, prediabetes-AAB^HPAP^, T2D^HPAP,^ and T2D^PFKFB3^.

To find out whether PFKFB3 is a gatekeeper of CFC in diabetes (Montemurro *et al*., 2019; Nomoto *et al*., 2020), we sought to establish whether PFKFB3 positive β-cells are identical to “loser” β-cells from CFC. To conceptualize our objective, we considered β-cells with different competitive traits depicted in **Fig 1C**. “Loser” β-cells (L) which don’t express PFKFB3 are elective for CFC. However, “loser” β-cells that express PFKFB3 (P+L) are non-elective by CFC because of PFKFB3 protection. Alone PFKFB3 (P) expression represents an indolent CFC status because it doesn’t co-occur with the “loser” signature (**Fig 1C**).

Gene-set enrichment analysis (GSEA) based on DAVID and using Reactome as a reference unraveled differential transcriptomes (p<0.05) of “loser” ϕ3-cells from Control^HPAP^, AAB^HPAP^, T1D^HPAP^, T2D^HPAP^ and T2D^PFKFB3^ (**Fig 1D, E**). DEGs submitted to GSEA were normalized with the union of all genes from the HPAP database or the union of all genes from the nPOD database, respectively (**Figs 1D, E,** and **Tables S1-S5**). The percentage coverage (ratio of the observed gene to background gene count) increased while the strength of enriched pathways decreased from prediabetes (AAB ^HPAP^) to T2D^HPAP^ with T2D^PFKFB3^ accounting for an intermediate state (**Fig 1D and Fig S1**). Pair-wise overlap between each two disease states is presented in **Fig S2A-C.**

Enriched pathways referred to the genes encoding for proteins involved in ribosomal biosynthesis (RiBi), protein translation, and peptide elongation processes in all datasets while enriched pathways of mitochondrial respiration dominated the T2D^HPAP^ dataset (**Fig EV2**). Among the RiBi-related enriched pathways in T2D^PFKFB3^, AAB^HPAP,^ and T2D^HPAP^ we found specifically *Protein translation, Peptide chain elongation, Selenoamino acid metabolism, Signaling by ROBO receptors, Nonsense Mediated Decay (NMD),* and many others, all presented in **Fig EV2** confirming alignment with the data in **Fig 1D**.

The DEGs from all datasets were dominated by downregulated genes (**Tables S1-S5**). Prior filtering by adjusted p-value, individual DEGs in AAB^HPAP^, T2D^PFKFB3,^ and T2D^HPAP^ showed global overlap in the direction of gene expression compared to Control^HPAP^ as demonstrated in the heatmap (scale represents log_2_FC) in **Fig EV3A, B**. We observed a subset of inversely correlated DEGs including regulators of insulin secretion such as the subunit of ϕ3-cell K^ATP^ channel *ABCC8*, phosphodiesterase *PDE8B*, a regulator of cAMP and cGMP degradation, bone morphogenic protein 5 (*BMP5*) which is involved in autophagy, *BAIAP3*, encoding for a Ca^2+^-dependent and RPH3AL Rab GTP effector of late exocytosis, the copper chaperone of superoxide dismutase (*CCS*) and *RGS16,* a component of the G-coupled receptor signaling, all of which were found upregulated in T2D^HPAP^ compared to T2D^PFKFB3^ (**Fig EV3B**). Folliculin (*FLCN*) was the only gene found upregulated in T2D^PFKFB3^ and downregulated in T2D^HPAP^ (**Fig EV3B**). FLCN can activate mTORC1 kinase by stimulating GTP hydrolysis of Rag GTPases (Ramirez Reyes *et al*, 2021). FLCN also plays a role in autophagy and modulation of glycolysis (Ramirez Reyes *et al*., 2021). Downregulation of ribosomal genes therefore constituted the major overlap in identical, individual DEGs (adjusted p value<0.05) from Control^HPAP^, AAB^HPAP^, T2D^HPAP,^ and T2D^PFKFB3^ (**Fig EV3C, D**).

We performed STRING analysis (Szklarczyk *et al*, 2021a, b) to identify protein-protein interactions based on physical evidence with high edge confidence (>90%) (Szklarczyk *et al*., 2021b) (**Fig 2A-C**). We found frameworks of protein associations with clusters visualizing the dominance of ribosomal genes (RiBi) in AAB^HPAP^ and T2D^PFKFB3^ and mitochondrial respiration with *Cytochrome-c oxidase activity*, *Ubiquinol-cytochrome-c reductase activity, Electron transfer activity, NADH-dehydrogenase (ubiquinone) activity* in T2D^HPAP^ dataset (**Fig 2A-C Fig EV4A-C**). Interestingly, in the mitochondrial respiration framework we identified one cluster related to *Mitochondrial Translation* and *ATP Synthases* both in T2D^HPAP^ and T2D^PFKFB3^, in the latter much less present (**Fig EV4A, B**). The connectivity framework of small and large ribosomal subunits was already present in prediabetes AAB^HPAP^ corroborating ribosomal genes as the origin of the ;-cell “loser” status. T2D^PFKFB3^ represented an intermediate state between prediabetes AAB^HPAP^ (where the mitochondrial respiration cluster was omitted) and T2D^HPAP^ (with the most prominent mitochondrial respiration cluster). These results suggested that the natural history of the “loser” ;-cell signature (“loser” trajectory) is associated with ribosomal biosynthesis which dominates early- and mitochondrial respiration dominating the late diabetes phenotype (**Fig 2A**, top right panel).

**Figure 2.**
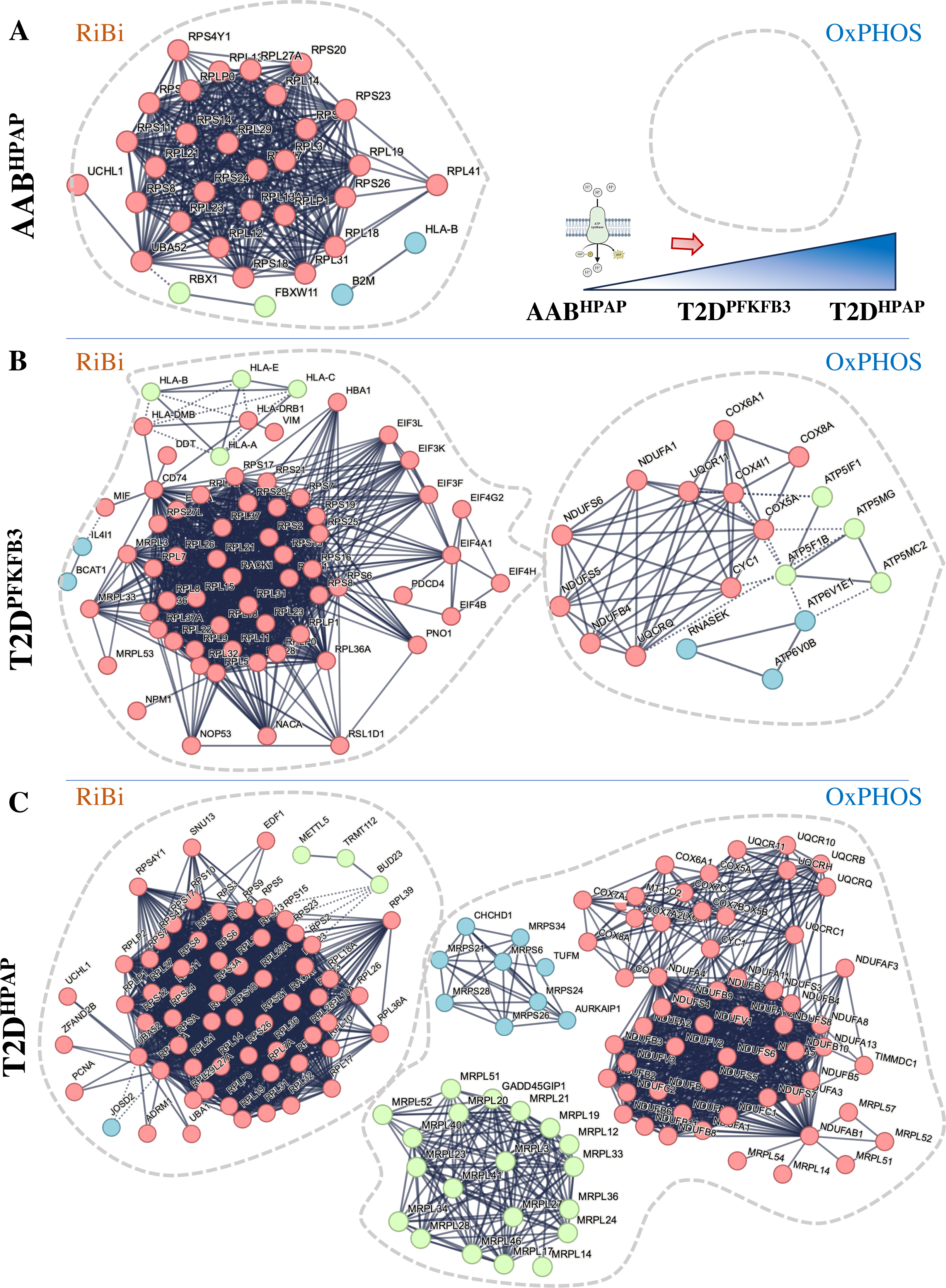
STRING analysis reveals ribosomes, mitoribosomes, and mitochondrial respiration are at the core of the “loser” signature. An experimental evidence (>90% confidence level) based protein-protein interaction framework of DEGs with a cluster of ribosomal biosynthesis and absence of the cluster for mitochondrial respiration in AAB^HPAP^ B Experimental evidence (>90% confidence level) based protein-protein interaction framework of DEGs with clusters for ribosomal biosynthesis and mitochondrial respiration in T2D^PFKFB3^ C Experimental evidence (>90% confidence level) based protein-protein interaction framework of DEGs with clusters for ribosomal biosynthesis and mitochondrial respiration in T2D^HPAP^ Ribosomal biosynthesis (RiBi), Oxidative phosphorylation (OxPHOS). Drawing in panel (A, top right) represents the “loser” ;-cell trajectory with downregulation of OxPHOS related genes emerging in PFKFB3 positive ;-cells and dominating T2D^HPAP^ but being absent in AAB^HPAP^.

Given that T2D^HPAP^ and T2D^PFKFB3^ may only differ relative to the active or inactive CFC (protected “loser” ;-cells), respectively, the unique clusters to T2D^PFKFB3^ may hold a clue to CFC inactivation. STRING analysis revealed downregulation of human leukocyte antigens (HLA) Class I (*HLA-A, HLA-B, HLA-C, HLA-E)* and Class II (*HLA-DRB1, HLA-DMB), Histones*, and components of the *Trypsinogen pathway* forming clusters that were specific only in T2D^PFKFB3^ (**Fig 2B, EV4B**). To corroborate these intriguing observations, we carried out an independent analysis.

### hdWGCNA and WGCNA reveal hub genes related to RiBi in all disease states

We applied the R package high dimension (hd)WGCNA (Morabito *et al*, 2023) on HPAP sc-RNA-seq data for each disease state. We obtained various networks as an output in the form of co-expression modules with the top 10 hub genes. Co-expression modules in each disease condition were correlated with “metacell” states of pancreatic β-cells (**Table S7-S10**). The “metacells” are defined as groups of transcriptionally identical and distinct cell states. They were created using a bootstrapped aggregation (bagging) algorithm by applying the K-nearest neighbors (KNN) method (Morabito *et al*., 2023).

To summarize the expression of the entire module into a single metric, module eigengenes (ME) were created for each metacell population. MEs are defined as the first principal component of the module’s gene expression matrix that describes the entire co-expression module. The turquoise co-expression module dominated because of the overt number of co-expressed genes in transcriptionally identical metacells for each disease condition (**Fig 3A-J**). This was corroborated by the falling dendrograms in which among the disease states the turquoise module was largest in T2D^PFKFB3^ (**Fig A, C, E**). The grey module comprised genes that were not grouped into any co-expression module and were, therefore, excluded from the subsequent analysis. To identify co-expression networks based on the strong gene-gene co-expression relationships without the noise of weak correlations, soft power thresholds of 12, 16, 20, and 14 were selected for mean, median, and max connectivity to reach the Scale Free Topology Model Fit greater than 0.8 for Control^HPAP^; AAB^HPAP^; T1D^HPAP^, and T2D^HPAP^, respectively (**Fig EV5)**. Feature plots of different co-expression modules are presented in **Fig S3**.

**Figure 3.**
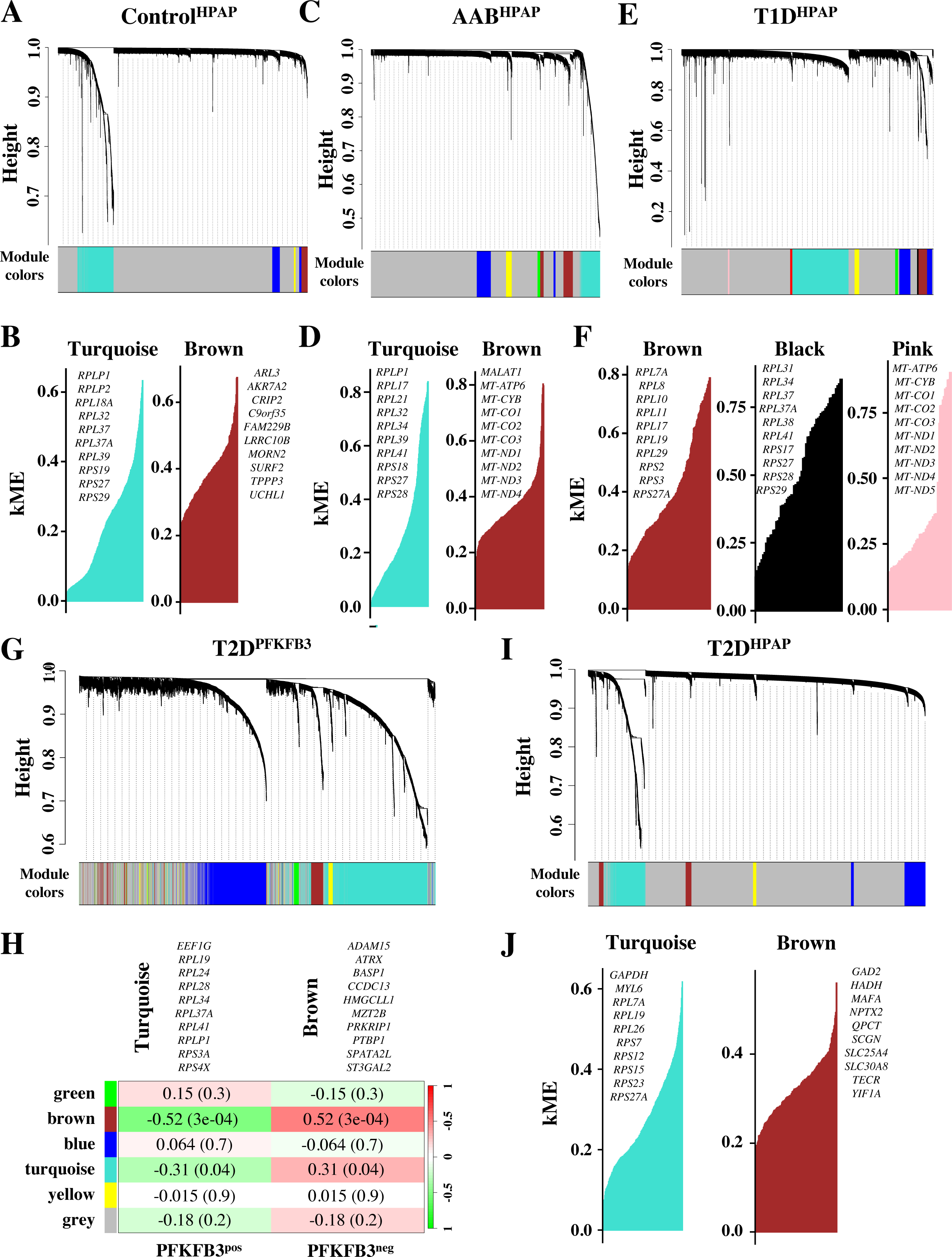
hdWGCNA and WGCNA reveal hub genes related to RiBi in all disease states. A-J Falling dendrograms and module eigengenes representing transcriptionally identical “metacell” states in Control^HPAP^ (A), AAB^HPAP^ (C), T1D^HPAP^ (E), PFKFB3 positive ;-cells i.e. T2D^PFKFB3^ (G), and “loser” ;-cells in T2D i.e. T2D^HPAP^ (I). The top 10 hub-genes with connectivity constant for module eigengenes (kME) of each hub gene showing dominant turquoise (RiBi) and brown (OxPHOS) modules for Control^HPAP^ (B), AAB^HPAP^ (D), dominant brown (RiBi), black (RiBi) and pink (OxPHOS) modules for T1D^HPAP^ (F), dominant turquoise (RiBi) and brown (OxPHOS) module eigengenes in T2D^PFKFB3^ with marked Z-score and p-values (H), and dominant turquoise (RiBi) module with kME value in T2D^HPAP^. Each leaf on the dendrogram represents a single gene, and the color at the bottom indicates the co-expression module assignment.

The eigengene-based connectivity (kME) with each gene in the scRNA-seq data assisted in revealing the hub genes of each module. In **Fig 3A** we presented only turquoise and brown modules out of four types of MEs (blue, turquoise, brown, and yellow) found in the Control^HPAP^ and they overlapped with the GSEA analysis. The turquoise module eigengenes comprised of RiBi-related hub genes such as RPLP1, RPL32, RPL37, RPL18A, RPS19, RPLP2, RPL37A, RPS29, RPS27, RPL39, constituting a shared feature between Control^HPAP^ and other datasets. In the AAB^HPAP^ dataset, we found turquoise and brown module eigengenes comprised of ribosomal genes and mitochondrial genes, respectively (**Fig 3C, D**).

Interestingly, in T1D^HPAP^ we found two module eigengenes (brown and black) related to ribosomal hub genes and pink module eigengenes for the mitochondrial hub genes. This contrasted the results referring to DEGs derived from “loser” criteria in T1D. Thus, T1D unlike other disease states may hold different “loser” criteria for selection of ribosomal and mitochondrial hub genes (**Fig 3E, F**). Surprisingly, mitochondrial modules were found in AAB^HPAP^ and T1D^HPAP^, different from enrichment analysis (**Fig 3D, F**). Since hdWGCNA is computed from the normalized gene expression matrix to generate metacell expression matrices using the KNN parameter, hub genes in MEs might not necessarily overlap with DEGs from the sc-RNA-seq analysis.

To be able to compare the MEs between T2D^PFKFB3^ and the T2D^HPAP^, we used conventional WGCNA (Langfelder & Horvath, 2008) for analysis of the matrix from spatial transcriptomics. We identified brown and turquoise modules in T2D^PFKFB3^ comprising genes involved in β-cell function (p<0.05) and ribosomal hub genes (p<0.05), respectively (**Fig 3G, H**). Similar to T2D^PFKFB3^, in the T2D^HPAP^ condition we found a dominating turquoise ME in reference to ribosomal subunits together with GAPDH and MYL6 but not the module eigengenes for the mitochondrial hub genes, which was different from the results of the GSEA of T2D^HPAP^ (**Fig 3I, J**). Thus, collectively, the turquoise module for ribosomal hub genes was shared across all data sets and was not correlated to the module with mitochondrial hub genes (**Fig S4**).

### Key transcriptional drivers of “loser” signature in β-cells

We used Mergeomics wKDA analysis to determine the key drivers of the differential gene expression in disease using gene network topology and edge weight information from DEGs (see Methods) (**Fig 4A-F**). Herein we identified networks of functionally related candidate hub genes with regulatory roles in the disease gene expression networks. In AAB^HPAP^ key drivers were represented by genes playing an important role in the elongation step of protein synthesis (*EEF2, EIF2S1, RPL26, RPL4,* and *RPLS1*) with *EEF2, RPL4,* and *RPLS1* making the functional connection **(Fig 4A, B**).

**Figure 4.**
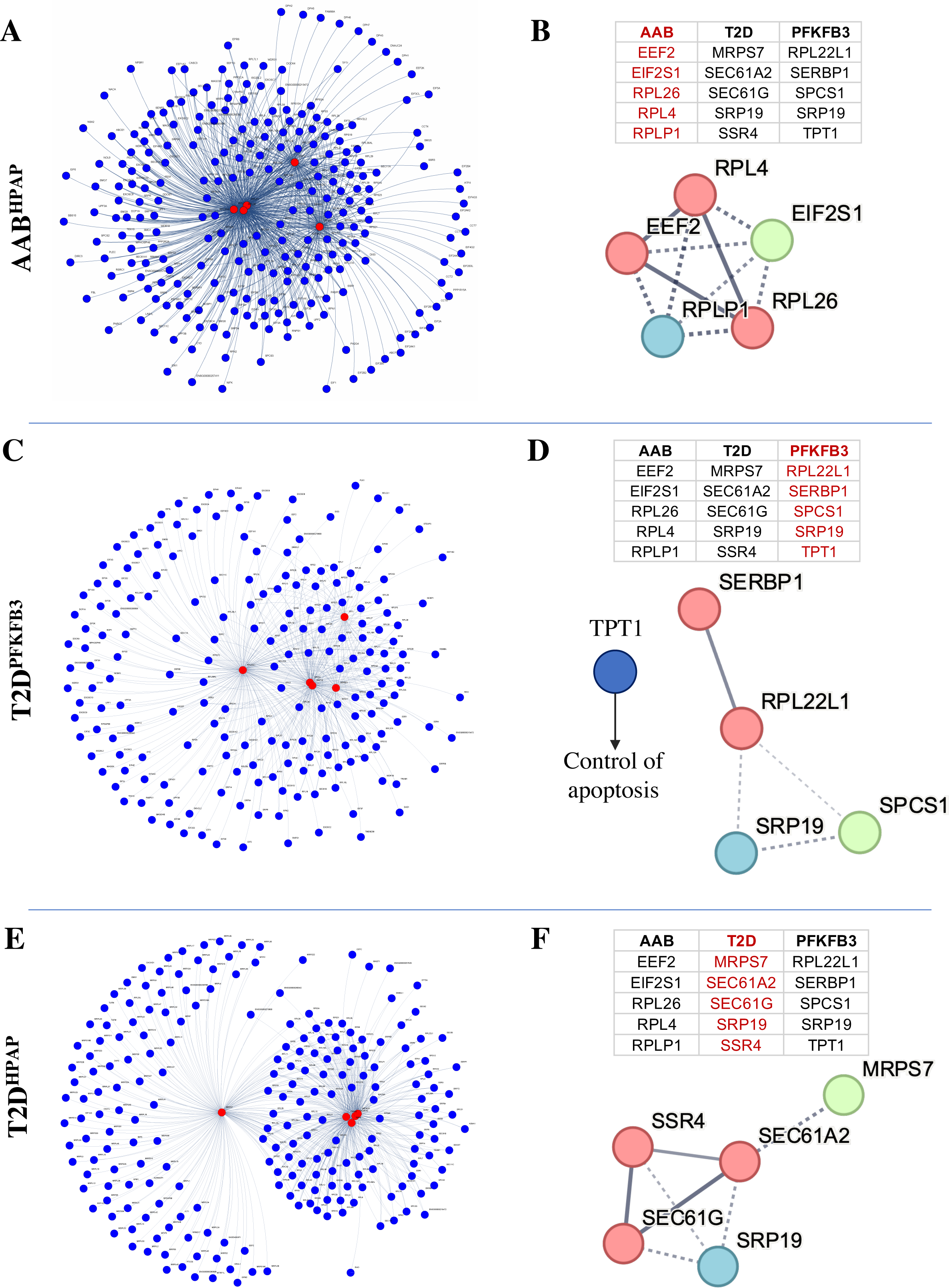
Key transcriptional drivers of “loser” signature in ;-cells. A, F Network of key-driver hub genes presented as red nodes and shared genes as blue nodes in AAB^HPAP^ (A), T2D^PFKFB3^ (C), and T2D^HPAP^ (E) with list of top five key driver genes (highlighted in red) in the adjacent table and the respective protein-protein interaction network in AAB^HPAP^ (B), T2D^PFKFB3^ (D), and T2D^HPAP^ (F). In T2D^PFKFB3^ TPT1 anti-apoptotic gene is not related to the cluster (D).

T2D^PFKFB3^ shared the *SRP19* key driver gene with T2D^HPAP^. Other key drivers in T2D^PFKFB3^ included Ribosomal Protein L22 Like 1 (*RPL22L1*), and the structural constituent of ribosome and SERPINE1 mRNA Binding Protein 1 (*SERBP1*) involved in SERPINE1/PAI1 mRNA stability and ribosome hibernation, a process during which ribosomes are prevented from proteasomal degradation (Shetty *et al*, 2023). We also identified Signal Peptidase Complex Subunit 1 (*SPCS1*) that catalyzes the cleavage of nascent proteins during their translocation in ER (Liaci *et al*, 2021), and Tumor Protein Translationally-Controlled 1 (TPT1) (Chen *et al*, 2013). Upon glucose stimulation, TPT1 is translocated to the mitochondria and the nucleus. Gene knockdown of *TPT1* induces apoptosis and its overexpression reduces ER-stress-related apoptosis (Diraison *et al*, 2011). (**Fig 4C, D**). Similar to TPT1, SERBP1-dependent fatty acid synthesis is required for insulin secretion at high glucose concentrations (Diraison *et al*, 2008). In addition, *RPL221*, *SERBP1,* and *TPT1* key drivers in T2D^PFKFB3^ not only formed functional connections but are all related to increased resistance to stress [*RPL22L1* and *SERBP1* in ribosome hibernation (Shetty *et al*., 2023), and *TPT1* in resistance to oxidative and ER stress (Chen *et al*., 2013)] (Diraison *et al*., 2011).

In T2D^HPAP^ key drivers were presented by Mitochondrial Ribosomal Protein S7 (*MRPS7*) from mitochondrial protein synthesis, SEC61 Translocon Subunit Alpha 2 (*SEC61A2*), a signal peptide-containing precursor for co-translational translocation of nascent polypeptides across the endoplasmic reticulum (ER), forming an ER ribosome receptor and a gated pore, Signal Sequence Receptor Subunit 4 (*SSR4*) that binds calcium and regulates retention of ER-resident proteins, SEC61 Translocon Subunit Gamma (*SEC61G*) that exerts ATPase activity, and Signal Recognition Particle 19 (*SRP19*) that enables 7S RNA binding activity. The functionally connected key drivers in T2D^HPAP^ comprised *SEC61* subunits and *SSR4*, falling under mitochondrial and ER regulatory genes (**Fig 4E, F**).

Collectively, these analyses confirmed that PFKFB3 positive β-cells are cells with “loser” signatures that show reduced HLA expression, evidence of potentially an escape from immunosurveillance. Since PFKFB3 expressing “loser” ;-cells are long-lived cells carried on from prediabetes, we asked about the mechanism by which PFKFB3 can exert protection despite the global distortion of RiBi and mitochondrial respiration.

### SNP-SNP analysis reveals PFKFB3 *rs1983890* interplay with anti-apoptotic gene *MAIP1*

The epiphenomenon of positive epistasis that can counter the global distortion of RiBi has been described in yeast (Tutaj *et al*, 2023) and we set to investigate it in relation to PFKFB3 gene-gene interactions (GGI). We analyzed the significant interactions between the PFKFB3 polymorphic allele on chromosome 10p15.1 locus (*rs1983890*) (Wallace *et al*, 2015) and all annotated SNPs from the HPAP dbGAP database corresponding to the list of DEGs between PFKFB3 positive and negative β-cells (**Fig 5A**). Using case (all diabetic donors) to control (non-diabetic donors) approach, we found only two genes with significant (adjusted p<0.05) SNP interactions, matrix-AAA peptidase interacting protein 1 (*MAIP1*) (adj. p value = 0.0113) and ArfGAP with GTPase domain, ankyrin repeat and PH domain 1 (*AGAP1*) (adj. p value = 0.0275) presented in the **Table S11 & S12**. MAIP1 is a novel mitochondrial matrix AAA-peptidase interacting protein 1, known for its interaction with the m-AAA peptidase, a component of the mitochondrial calcium uniporter (MCU) involved in the degradation of essential MCU regulator (EMRE).

**Figure 5.**
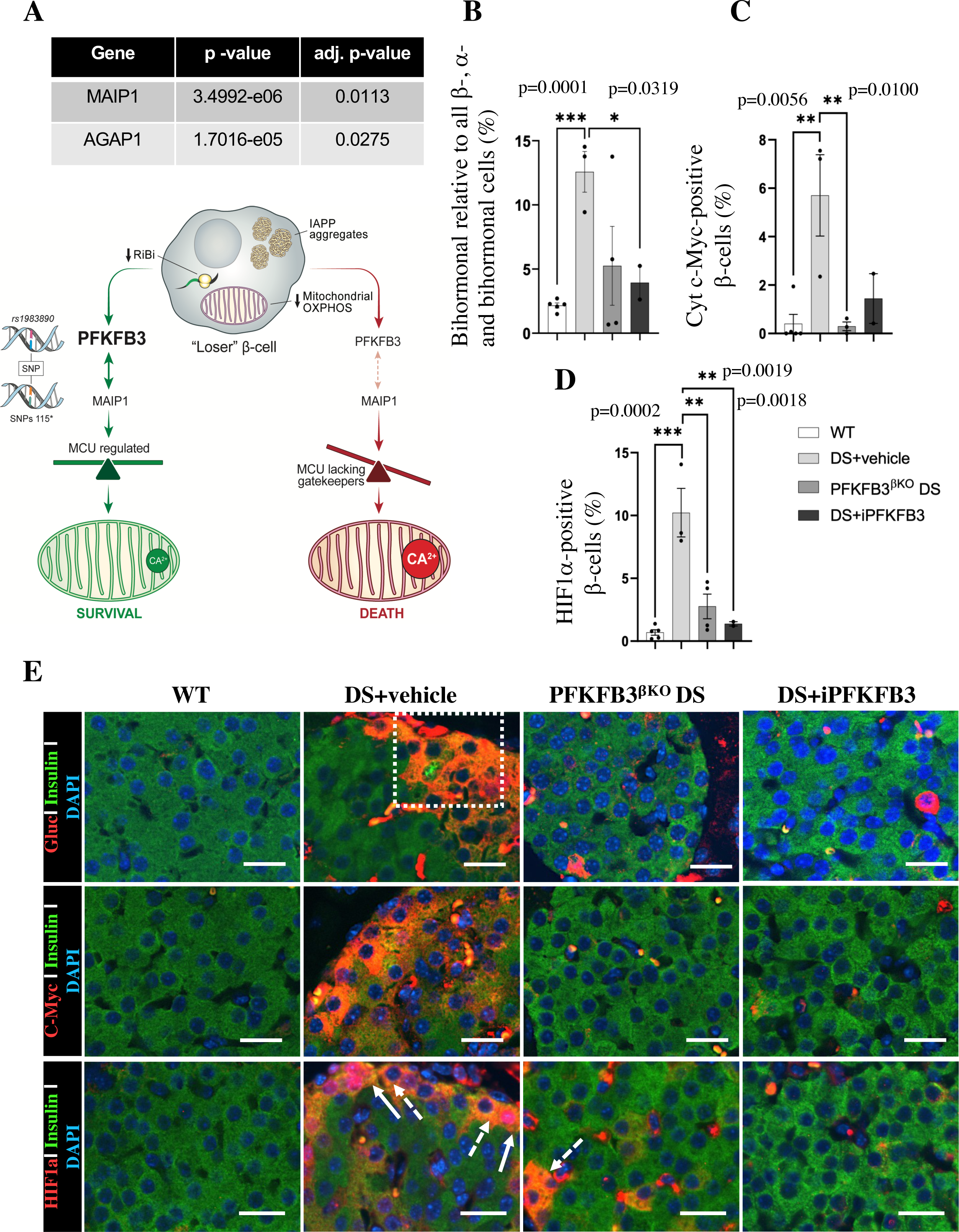
PFKFB3 inhibition eliminates dysfunctional ;-cells that are protected by positive epistasis. A PFKFB3 rs1983890-SNP interaction analysis with the DEGs in PFKFB3 positive relative to PFKFB3 negative ;-cells revealed by spatial transcriptomics yielded significant interactions with collective SNPs of AGAP1 and MAIP1 (SNPs 115). Diagrammatic reconstruction of the results indicates a mechanism of PFKFB3-mediated survival of injured “loser” ;-cells. MAIP1 is an anti-apoptotic gene that controls mitochondrial Ca2+ via MCU uniporter. MCU is composed of channel-forming subunits, EMRE, and a gatekeeper subunit MICU. The gating of the MCU complex requires an association between EMRE and MICU. MAIP1 may protect EMRE from i-AAA mediated degradation (28) preventing constitutive activation of MCU and cell death. B Quantitative representation of insulin-positive and glucagon-positive (bihormonal) injured cells (% of all β-, α and bihormonal cells) C Quantitative representation of HIF1α-positive injured β-cells (% of all ;-cells) D Quantitative representation of c-Myc positive injured ;-cells (% of all ;-cells) E Representative immunofluorescence images of islets, from wild-type mice exposed to high-fat diet, HFD (WT), mice under diabetogenic stress treated with vehicle (IAPP+HFD= DS; DS + vehicle), mice with conditional PFKFB3 knockout exposed to diabetogenic stress for 6 weeks (PFKFB3^βKO^ DS), and mice under diabetogenic stress receiving PFKFB3 inhibitor AZ67 for 6 weeks (DS + iPFKFB3), either double immunostained for insulin- and glucagon (to reveal bihormonal cells), or immunostained for cytoplasmic c-Myc, and HIF1α (all markers in red), insulin (green) and nuclei (blue) showing eradication of marker positive injured ;-cells in PFKFB3^βKO^ DS and DS + iPFKFB3 mice. Scale bar: 25 µm.

These results indicated that PFKFB3 may play a direct role in the survival and protection of “loser” ;-cells, indicating that targeting PFKFB3 can facilitate the clearance of these cells.

Therefore, we set out to analyze the phenotypic consequences of PFKFB3 depletion or inhibition in mice undergoing diabetogenic stress and the feasibility of targeting PFKFB3 at a systemic level.

Imaging analysis of the immunostainings from the hβTG mouse pancreata from compared to the wild-type (WT) controls, indicated accumulation of different types of injured β-cells such as double insulin- and glucagon (bihormonal), cytoplasmic c-Myc positive-, and HIF1α positive β-cells (**Fig 5B-E**). We haven’t found a transcriptional upregulation of PFKFB3 in “loser” β-cells. However, PFKFB3 could be stabilized in a post-translational fashion implicating APC/Cdh1 and Emi1 (Cappell *et al*, 2018; Tudzarova *et al*, 2011). In that sense we found a downregulation of APC/Cdh1 (*FZR1*) in T2D^HPAP^ (log_2_FC=-0.21; p=0.004) whereas in T2D^PFKFB3^ downregulation was not significant (log_2_FC=-0.004; p=0.6) (**Table S13**).

PFKFB3 expressing β-cells exceed the number of injured β-cells and PFKFB3 knockout leads to clearance of all injured β-cells (Min *et al*., 2022). We separated the PFKFB3 targeting group (n=5) into 3 animals subjugated to β-cell-specific knockout of PFKFB3 (PFKFB3^βKO^ DS) as a positive control (Min *et al*., 2022) and two animals that received PFKFB3 inhibitor AZ67. PFKFB3^βKO^ DS led expectedly to the diminishment of all types of injured β-cells (**Fig 5B-E and S5-7**). 7 weeks of i.p. administration of the PFKFB3 inhibitor AZ67 (DS+ iPFKFB3 group) recapitulated the effect of PFKFB3^βKO^ DS as confirmed with quantification of all injury markers positive-versus all β-cells (**Fig 5B-E**). Bihormonal cells (that are increased in HFD), HIF1α positive - (dysfunctional β-cells reflecting islet inflammation induced by HFD), and cytoplasmic c-Myc positive β-cells (indicators of cells undergoing hIAPP-induced calpain activation and toxicity) which were significantly upregulated in DS+vehicle, were depleted in DS+iPFKFB3 mice (**Fig 5B-E**).

We asked whether clearance of injured “loser” β-cells under DS conditions would be sufficient to restore metabolic control in the DS mice.

### Systemic PFKFB3 inhibition (iPFKFB3) improves metabolic performance of hβTG (T2D) mice

We administered PFKFB3 inhibitor AZ67 daily (at 28 mg/kg body weight) from day one of exposure to HFD for 7 weeks. We measured IP-GTT at 4 and 6 weeks of iPFKFB3 treatment and ITT at 5 and 7 weeks of iPFKFB3 treatment. The experimental timeline of PFKFB3 inhibitor (iPFKFB3) administration and metabolic measurements are depicted in **Fig 6A**.

**Figure 6.**
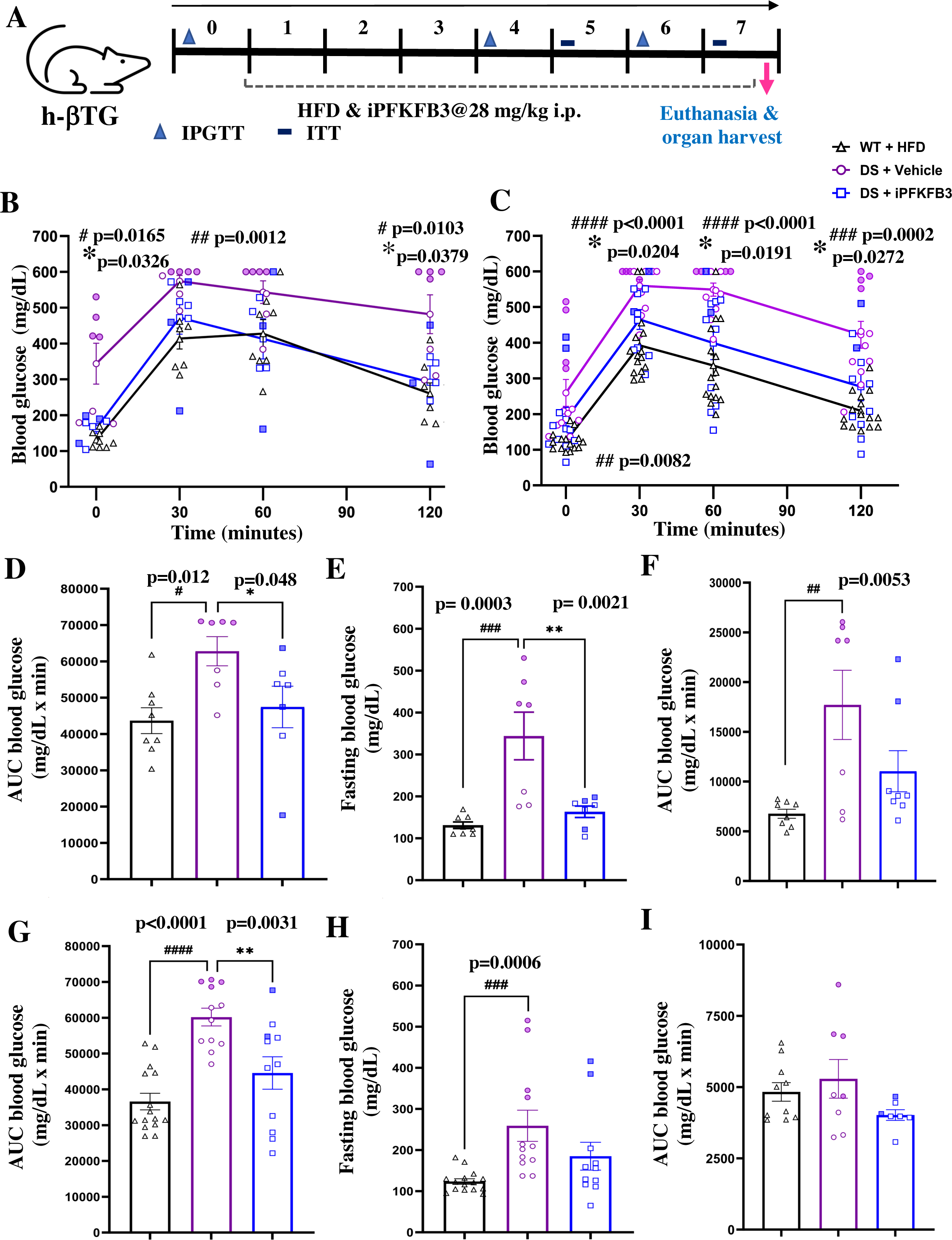
Systemic PFKFB3 inhibition (iPFKFB3) improves metabolic performance of h;TG (T2D) mice. An Experimental scheme of PFKFB3 inhibitor treatment of h;TG mouse; PFKFB3 inhibitor was administered at 28 mg/kg dose via intraperitoneal route for 7 weeks and metabolic tests were performed (IPGTT and ITT) at indicated time points. B, C Intraperitoneal glucose tolerance test (IP-GTT) profile at 4 weeks (B) and 6 weeks (C) showed significant improvement in glucose tolerance in the DS + iPFKFB3 as compared to DS + vehicle mice. D-I Comparison of the area under the curve of IP-GTT (D) and fasting blood glucose (E) at 4 weeks and 6 weeks (G, H) showing significant improvement in DS + iPFKFB3 as compared to DS + vehicle mice; Area under the curve (AUC) of insulin tolerance test (ITT) at 5 weeks (F) and 7 weeks (I) showing a trend of improvement in insulin sensitivity in DS + iPFKFB3 mice as compared to DS + vehicle mice without reaching significance 4 weeks: n=8 or 7, n=7 and n=7 biological replicates for WT+HFD, DS + vehicle and DS + iPFKFB3, respectively; **#** WT vs DS+vehicle, **\***DS+iPFKFB3 vs DS+vehicle. 6 weeks: n=15, n=12 and n=11 biological replicates from n=2 independent experiments for WT+HFD, DS + vehicle and DS + iPFKFB3, respectively. Data is represented as Mean ±SEM, p<0.05. Full coloured graph symbols (circle, triangle and square) represent mice where hIAPP transgenic signal as obtained from genotyping was more than 8 arbitrary units in reference to higher hIAPP expression and proteotoxicity. Hollow symbols correspond to mice with hIAPP transgenic signal higher than 4 and less than 8 arbitrary units.

The clearance of injured β-cells by PFKFB3 inhibition correlated with the improved glucose tolerance as demonstrated by IP-GTT and by AUC of blood glucose (±SEM, p<0.05) after 4 weeks (n=7-8, ±SEM, p<0.05) **(Figs. 6B and 6D, E)** and 6 weeks (2 independent experiments, n=4-8/group, ±SEM, p<0.05) **(Figs. 6C and 6G, H)**. Improved glucose tolerance led to reduced fasting blood glucose (**Fig 6E, H**). Moreover, insulin sensitivity was increased although not reaching significance in DS+iPFKFB3 mice after 5 weeks (n=7-8, ±SEM, p<0.05) **(Fig 6F)** and 7 weeks (2 independent experiments, n=4-8/group, ±SEM, p<0.05) **(Fig 6I)** when compared to DS+vehicle group.

Collectively, these data indicated that targeting PFKFB3 positive β-cells with “loser” signature by PFKFB3 inhibition leads to diminishment of dysfunctional “loser” β-cells, restoring glucose tolerance in human-like model of T2D.

## DISCUSSION

CFC has been successfully validated in post-mitotic cells (Coelho & Moreno, 2019; Coelho *et al*., 2018; Gradeci *et al*, 2021; Vieira *et al*., 2023) and here we evaluated the hypothesis that “loser” and “winner” cell imbalance may underly one of the pathogenic mechanisms in T2D.

We demonstrated that PFKFB3 positive β-cells are “loser” cells and that targeting PFKFB3 can facilitate “loser” β-cell clearance by re-activation of CFC. β-cell “loser” signature emerged from the high level of transcriptional convergence between AAB^HPAP^, T2D^HPAP,^ and T2D^PFKFB3^ based on dual criteria to reveal “loser” β-cells. One criterion included the UPR and oxidative stress-related genes that confer “loser status” (Baumgartner *et al*., 2021) and are upregulated in mouse epiblast “loser” cells (Lima *et al*., 2021). The second criterion was based on PFKFB3 expression that is triggered by IAPP proteotoxicity and cytokine stress in T2D and T1D, respectively (Min *et al*., 2022; Montemurro *et al*., 2019). We established the similarities and differences comparing the DEGs between “loser” and “non-loser” ;-cells across 4 disease states (Control, AAB, T1D, and T2D) from the HPAP scRNA-seq and from spatial transcriptomics on nPOD T2D pancreata to distinguish PFKFB3 positive- and negative β-cells. Although we have not found transcriptional upregulation of PFKFB3 in the “loser” HPAP datasets, we found that APC/Cdh1 E3 ubiquitin ligase that regulates PFKFB3 stability (Tudzarova *et al*., 2011) was downregulated in T2D^HPAP^. These observations indicate that “loser” β-cells in T2D could stabilize PFKFB3 post-translationally, as previously suggested (Wigger *et al*, 2021).

Both RiBi and mitochondrial genes were enriched in the DEGs and metacells from WGCNA, respectively, although there was no correlation between these two in WGCNA. These results indicated that the “loser” β-cell signature based on RiBi potentially represents a measurable metacell entity across all β-cells in all disease states and that it evolves from prediabetes to sincere T2D. PFKFB3 positive β-cells in T2D shared the signature of global stochiometric distortion of RiBi with “loser” β-cells in AAB^HPAP^ and T2D^HPAP^. These findings are interesting since originally the “loser” status was demonstrated by a dominant ribosomal subunit (Minute) mutation in *Drosophila* (Carnegie Institution of, 1923). The enrichment showing distortion of RiBi in Control^HPAP^ hasn’t reached significance, which may be due to an active CFC in health. While the overlap existed between β-cells from AAB^HPAP^, T2D^HPAP^, and T2D^PFKFB3^, there was no shared signature with T1D. Interestingly, metacells in T1D were represented by both ribosomal and mitochondrial hub genes thereby not correlating with GSEA and DEGs based on UPR and oxidative stress criteria. This lack of correlation may point to different CFC mechanism(s) and/or enhanced attrition of “loser” β-cells by autoimmunity in T1D. T2D^HPAP^ was dominated by the downregulation of mitochondrial respiration genes in GSEA. T2D^PFKFB3^ pertained to the global shutdown of RiBi, had yet only one enriched category of genes encoding mitochondrial respiration, positioning itself between AAB^HPAP^ and T2D^HPAP^. The transcriptional proximity of AAB^HPAP^ and T2D^PFKFB3^ was also evident in the subset of the overlapping genes presented in the heatmaps (**Fig EV3A, C-D**). Interestingly, the emerging difference in the order from AAB^HPAP^ to T2D^PFKFB3^ and T2D^HPAP^ can be highlighted by the downregulation of the ribosomal gene encoding RPS12 in T2D^HPAP^ and T2D^PFKFB3^. RPS12 is required for *Xrp1* [human *DDIT3* (Blanco *et al*, 2020)]-dependent regulation of protein translation, growth, and CFC (Ji *et al*, 2019). Since a missense mutation in RPS12 can protect “loser” cells from attrition (Kale *et al*, 2018), the observed downregulation of *RPS12* in our studies may be in line with increased survival of “loser” β-cells in T2D^HPAP^ and T2D^PFKFB3^.

The profound downregulation of RiBi and genes of mitochondrial respiration in “loser” β-cells indicate that conserving energy by shutting down the energy-expensive process of making proteins is a priority of long-lived dysfunctional ;-cells. That was confirmed by the enrichment of GO terms in PFKFB3 positive “loser” β-cells that pertain to mRNA and protein quality control during translation at the ribosome and/or ribosome hibernation (*RPL22L1, SERBP1, and TPT1*). We observed strong transcriptional downregulation of cap-dependent and cap-independent translational machinery that triggers initiation via internal ribosomal entry sites (IRES). Stalling of the initiation step is accomplished via post-translational modification of eIF2a (Richter & Coller, 2015) that is complemented by transcriptional suppression as we observed here. Further studies will unravel whether the “loser” signature in PFKFB3-positive ;-cells was originally introduced as a transient attempt to adaptively preserve some orphaned ribosomal proteins after RiBi shutdown (Buchan & Parker, 2009; Wheeler *et al*, 2016).

Since β-cells with PFKFB3 expression are detected decades after onset of T2D, we reasoned that breaking down their unique genetic makeup that differs from T2D^HPAP^ might offer a clue to the long-term survival of PFKFB3 positive “loser” β-cells. One of the most remarkable differences in T2D^PFKFB3^ was the downregulation of human leukocyte antigen (HLA) class I and II implicated in immune surveillance (Adachi *et al*, 2022; Drake *et al*, 2006). This is in line with the reduced haplotype frequency of HLA related to *PFKFB3* polymorphism (Chen & Chen, 2019). Downregulation of HLA is connected to a fail-safe response by natural killer (NK) cells (Shi *et al*, 2011), however chronic inflammation that normally co-occurs with diabetes may suppress NK cells allowing PFKFB3 “loser” ;-cells to bypass the immunosurveillance. The unique HLA makeup in T2D^PFKFB3^ poses an intriguing possibility that immunity plays a role in the removal of PFKFB3-positive “loser” β-cells (Johnston, 2009), which will be addressed in the future.

Another mechanism to explain the prolonged survival of the PFKFB3 positive “loser” β-cells may involve epistasis. Epistasis is a major contributor to variations in disease outcome and it refers to the dependence of a mutation on other mutation(s) and the genetic context in general. Epistasis was demonstrated in British and French pedigrees involving T2D susceptibility loci on chromosomes 1q21-25 and 10q23-26 (Cordell, 2002). The new PFKFB3 single nucleotide polymorphism (SNP) at the 10p15.1 locus (*rs1983890*) (Wallace *et al*., 2015) was predicted to exert genome-wide significance in diabetes (Chen & Chen, 2019). Our GGI analysis revealed PFKFB3 SNP interaction with *MAIP1* and *AGAP1* collective SNPs (Fig 5A). Disruption of *AGAP1* function can chronically activate the integrated stress response (IRS), leaving AGAP1-deficient cells susceptible to a variety of stressors (Lewis *et al*, 2023). On the other side, MAIP1 (C2ORF47) was first identified via the neuronal interactome of *m*-AAA proteases in mice, and it was found to represent an anti-apoptotic gene that regulates mitochondrial ribosomal trafficking and the assembly of the mitochondrial Ca^2+^ uniporter MCU. Thus, MAIP1 may prevent mitochondrial Ca^2+^ overload, mitochondrial permeability transition pore opening, and cell death (Konig *et al*, 2016). Interaction of PFKFB3 *rs1983890* with *MAIP1* and *AGAP1* SNPs may explain the survival of dysfunctional β-cells despite heightened ISR and deregulated mitochondrial Ca^2+^ homeostasis in both T2D and T1D (Huang *et al*, 2010).

CFC provides a context-dependent removal of injured or dysfunctional cells that has been proven successful through concepts of “synthetic lethality” and immune-checkpoint inhibition in the past (Lucchesi, 1968; O’Neil *et al*, 2017; Shiravand *et al*, 2022). To unlock CFC and demonstrate feasibility at the systemic level of PFKFB3 targeting, we utilized small molecule PFKFB3 inhibitor AZ67 (Boyd *et al*, 2015). We successfully eliminated injured β-cells recapitulating PFKFB3^βKO^ DS, which led to improvement in glucose tolerance. Thus, PFKFB3 targeting can reactivate CFC and induce functional β-cell regeneration through islet replenishment with healthy functional β-cells.

The main limitation of the study arises from the fact that CFC is a temporal event that is linked to a silent phenotype (no change in β-cell mass) so only a study design that will involve imaging human islets transplanted in the eye chamber of NOD/SCID mice and used to monitor CFC in real time would ultimately capture CFC dynamic.

Collectively our results describe that the ribosomal biosynthesis and mitochondrial respiration (cellular energy conservation) represent a ‘global’ phenotypic interface of ;-cell fitness that in the presence of PFKFB3 can create an epiphenomenon of positive epistasis via *AGAP1* and *MAIP1*. Reduced HLA expression may help these cells to bypass immunosurveillance. Positive epistasis is necessary to suppress the competition between “loser” and healthy “winner” ;-cells favoring the survival and accumulation of dysfunctional ;-cells. Therefore, pharmacological targeting of PFKFB3 holds a promise to change the T2D trajectory at the early onset given that at this stage both the prevalence of “loser” ;-cells and “loser” ;-cell survival selectively depend on PFKFB3.

## MATERIALS AND METHODS

### Study design

We analyzed large-scale transcriptomic data of human pancreatic β-cells from two independent datasets. We used the Human Pancreas Analysis Program (HPAP) (Kaestner *et al*, 2019). HPAP is part of the Human Islet Research Network supported by the National Institute of Diabetes and Digestive and Kidney Diseases (NIDDK) which leverages deep phenotyping of the human endocrine pancreas, thereby accumulating, analyzing, and distributing high-value data sets to the diabetes research community through the HPAP-PANC-DB database. We also used formalin-fixed and paraffin-embedded pancreata from three T2D donors from the Network of Pancreatic Organ Donors with Diabetes program (nPOD) (Campbell-Thompson *et al*, 2012).

We analyzed scRNA-seq from 31 non-diabetics (Control^HPAP^), 10 prediabetics with or without dysglycemia classified based on 2 or more autoantibodies (AAB^HPAP^), 9 donors with Type-1 diabetes (T1D^HPAP^), and 17 donors with Type-2 diabetes (T2D^HPAP^) using HPAP annotation of pancreatic β-cells. For each condition, and based on the module score reflecting the expression (Log_2_FC >0.1) or not of the “loser genes” *DDIT3, Atf3, Ppp1r15a, RICTOR, and Nfe2l2* (Log_2_FC< or = 0), cells were split into *“loser signature”*-positive and *“loser signature”*-negative β-cells for differential expression analysis between the two groups (Ortiz-Barahona *et al*, 2010; Valvona *et al*, 2016). “Loser” genes were adopted from *bona fide* “loser signatures” established in mouse embryos in Lima et al. (2021) (Lima *et al*., 2021).

### Data processing and clustering with module score

We analyzed the scRNA-seq data in R (v4.3.1) (URL https://www.R-project.org/) using Seurat (v4.3.0.1) (https://www.rdocumentation.org/packages/Seurat/versions/4.3.0.1) (Hao *et al*, 2021), beginning with the PercentageFeatureSet function to identify and filter β-cells based on HPAP annotation. A total of 44,559 β-cells were analyzed after being classified as either “Controls” (27,160 cells), “AAB” (8523 cells), “T1D” (715 cells) and “T2D” (8161 cells). The NormalizeData function was then used to perform log-normalization of the data subsample. The FindVariableFeatures function was utilized to calculate gene variances as well as feature variances of standard and clipped values. A variable called “genes.of.interest/loser signature” was created in the module score and candidate marker genes that represent key determinants of “loser” status implicated in the unfolded protein response (*DDIT3, ATF3, PPP1R15A, RICTOR)* and oxidative stress *NFE2L2* were then assigned to this variable. The ScaleData function was used to center and scale the data matrix. The AddModuleScore function was used to assign module scores to our subsample of either Control, AAB, T1D, or T2D β-cells based on the candidate genes specified in the “genes.of.interest/loser signature”. Two matrices were then created: one matrix containing 0 and negative module scores, and the other matrix containing positive module scores. The FindMarkers function was used to find differentially expressed genes within each disease state subsample by comparing the positive module scores’ matrix to the matrix containing 0 or negative module scores. Post-clustering doublet removal was integral to the HPAP data quality control.

### GeoMx spatial transcriptomics

A geospatial technology platform (GeoMx from NanoString Technologies, Inc, Seattle, WA) was used to perform targeted transcriptomic profiling of β-cell populations from three pancreata donors from the nPOD collection based upon PFKFB3 expression at the Molecular Pathology at City-of-Hope National Center, Duarte, California. We used a ssDNA labeled probe with photocleavable indexing oligos to bind targets of interest in nPOD pancreatic sections. We also used PFKFB3- and insulin antibodies conjugated with fluorophores as morphology markers to visualize PFKFB3 positive β-cells. Regions of interest (ROI) with PFKFB3 positive- and negative β-cells within a single ROI (> 100 cells) were segmented and collected separately from each slide from 3 T2D donors (nPOD #6186, #6255, and #6275). We analyzed 12-16 ROIs from each T2D formalin-fixed pancreatic section (8 PFKFB3 positive and 8 PFKFB3 negative ROIs). ROI were sequentially exposed to UV light to release the indexing oligos, which were collected and subsequently enumerated via Next-generation sequencing. We performed differential profiling of PFKFB3 positive relative to PFKFB3 negative β-cells using GeoMx DSP software for Whole Transcriptome Atlas (WTA).

### Gene enrichment analysis

To identify gene expression modules that showed clear, interpretable enrichment for biological functions, we performed a gene enrichment analysis using DAVID (Huang da *et al*, 2009; Sherman *et al*, 2022) based on Reactome annotations which contained multiple modules with significantly enriched gene ontology profiles (adjusted p<0.05).

To avoid a potential bias resulting from differences between reference cells (β-cells) from all cells, the sets of differentially expressed genes (FDR < 5%) in Control, AAB, T1D, and T2D β-cells were analyzed with DAVID (Huang da *et al*., 2009; Sherman *et al*., 2022), using the union of genes observed in all the samples as the reference. Enriched Reactome pathways identified in the individual datasets (FDR <5%) were visualized with Cytoscape (Shannon *et al*, 2003).

### Cell-specific gene network analysis

We analyzed two independent datasets, β-cell populations from three pancreata donors from the nPOD collection based upon PFKFB3 expression in spatial transcriptomics and sc-RNA-seq data under four conditions from a β-cell population from the PANC-DB Data portal of the HPAP (Kaestner *et al*., 2019; Montemurro *et al*., 2019). Weighted Gene Correlation Network Analysis (WCGNA) (Langfelder & Horvath, 2008) was used to construct correlated gene modules. Sample quality control was performed with the WGCNA package function, goodSampleGenes. The soft power threshold was adjusted for each condition to reach scale-free topology. Highly correlated modules among different blocks were then merged to create the final network. Subsequent analysis was focused on the significantly different modules with criteria, p < 0.05. The top 10 genes for each significant module were highlighted. Single-cell WGCNA was performed using high dimension (hd)WGCNA (Morabito *et al*., 2023), with the principal component analysis used for dimensionality reduction, the maximum number of cells shared between two metacells limited to 10, single-cell transform of the expression data, and soft power threshold adjusted to reach scale-free topology (12 for the ;-cell subset) as a signed networkType with different values for the individual i.e. Control, AAB, T1D, and T2D conditions. The top 10 hub genes for each module were highlighted. To reduce the noise of the correlations in the adjacency matrix, soft-power thresholds of 12, 16, 20, and 14 were picked for mean, median, and max connectivity to reach the Scale Free Topology Model Fit greater than 0.8 for Control^HPAP^; AAB^HPAP^; T1D^HPAP^, and T2D^HPAP^, respectively of hub-genes in the hdWGCNA.

Gene expression analysis was performed using R (v4.3.1) (https://www.R-project.org/). Network visualization was performed with Mathematica® (v12, Wolfram Research, Inc., Champaign, IL) using GravityEmbedding for the graph layout.

### Mergeomics

We performed weighted Key Driver Analysis (wKDA) (Arneson *et al*, 2016; Ding *et al*, 2021) on the subsets of DEGs from geospatial targeted transcriptomic profiling of β-cell populations from three nPOD pancreata donors and from “loser signature” transcriptome profiling of HPAP sc-RNAseq on the University of California, Los Angeles Web server for multidimensional Data integration (http://mergeomics.research.idre.ucla.edu/home.php<u>#)</u>. We used the Mergeomics 2.0 Web server for multiomics data integration (Ding *et al*., 2021).

### STRING Analysis

STRING analysis was performed on the STRING Web portal https://string-db.org using DEGs from the two independent data sets (HPAP and nPOD) (Szklarczyk *et al*., 2021a, b).

### Gene-gene interaction (GGI) analysis

Whole genome sequencing (WGS) data from the Human Pancreas Analysis Program (HPAP) was downloaded from the NIH database of Genotypes and Phenotypes (dbGAP). Exonic variants were annotated and filtered for genes of interest using bcftools. We used bcftools version 1.11, and as reference genome we used the Genome Reference Consortium Human Build 39 patch release 13 (GRCh 38.p13 or hg38). All variants for the gene PFKFB3 were filtered out except for the variant of interest, *rs1983890*. To detect all possible pairwise interactions between *rs1983890* and other genes, we used GeneGeneInteR R package and employed the Principal Component Analysis (PCA) method. For this method, a likelihood ratio test is conducted to compare two generalized linear models, M_inter_ and M_0_, where M_inter_ includes an interaction term and is defined as:

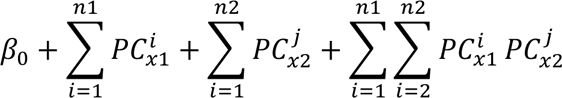

and

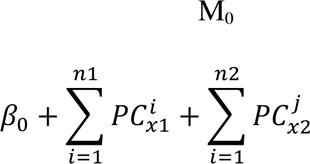

For both models, 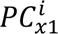 and refers to the *i*^th^ principal component of all SNPs in gene 1, 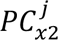 as the *j*^th^ principal component of all SNPs in gene 2. The number of principal components retained for each model, *n1* and *n2*, is found by the percentage of inertia calculated by PCA.

Following interaction testing, the resulting p-values were adjusted for multiple testing to obtain false discovery rate (FDR) using the Benjamini-Hochberg procedure.

### Animals

The study received ethical approval ARC-2019-011-AM-004 by the UCLA Animal Research Committee.

The β-cell-specific inducible PFKFB3-knockout or PFKFB3-wild type (WT) mouse model (*RIP-CreERT:PFKFB3^fl/fl^*) on *hIAPP^+/-^* background (h;TG) and high-fat diet (HFD) (PFKFB3^;KO^ DS) (#D12492, Research Diets Inc, New Brunswick, NJ, USA) was previously described (Min *et al*., 2022) and used as a positive control. At 6-8 weeks of age, all mice from the WT or DS group (h;TG) were assigned to receive the HFD. Additionally, mice from the DS group were randomly assigned to a group that received vehicle (DS+vehicle) or the group that received PFKFB3 inhibitor AZ67 (DS+iPFKFB3) (Tocris Biosciences). For the comparative evaluation of the effect of the PFKFB3 knockdown with the effect of PFKFB3 inhibition, we used AZ67 prepared in 10% absolute ethanol in sunflower oil, matched to the preparation of tamoxifen for the Cre-recombinase induction and PFKFB3 knockout. We randomized the PFKFB3 targeting group into mice (n=3) that received tamoxifen injection to undergo β-cell specific PFKFB3 knockout and mice (n=2) that received AZ67. For the rest of the experiments (two biological replicate experiments; each n=4-8/group), we used AZ67 prepared in 8% wt/vol 2-hydroxypropyl beta-cyclodextrin in PBS pH=7.4 at 28 mg/kg body weight intraperitoneally every day for 7 weeks **(Fig 6A)**. The mice had *ad libitum* access to food and water for the duration of the study. Body weights and fasting blood glucose levels were assessed weekly.

### Intraperitoneal glucose tolerance tests (IP-GTT)

An intraperitoneal glucose tolerance test (IP-GTT) was performed at 4 and 6 weeks after the start of the HFD and AZ67 administration. Mice were fasted overnight in a clean cage and with access to water before the analysis. Tail vein blood glucose was collected before and 30, 60, and 120 minutes after an injection of 20% glucose bolus (2 g/kg of body weight). Fasting blood glucose was measured weekly after overnight fasting for 15 hours. The blood glucose levels were measured in tail-drawn blood through the use of a freestyle blood-glucometer (Abbott Diabetes Care Inc, Alameda, CA, USA).

### Pancreas perfusion and isolation

Mice were euthanized by a brief isoflurane exposure before cervical dislocation. A medial cut was made to open the abdomen and chest cavities. A cut of the right ventricle was followed by a poke of the left ventricle with a needle to inject 10 ml of cold phosphate-buffered saline (PBS) slowly for perfusion of the pancreas. After perfusion, the pancreas was placed in cold PBS and separated from other tissues, including the surrounding fat. The pancreas was then weighed before and after the excess PBS had been absorbed into lint-free tissue.

### Histological assessments

After the excision of smaller pieces, the pancreas was fixed overnight at 4 °C in 4% paraformaldehyde (Electron Microscopy Sciences 19202, Hatfield, PA, USA). The pancreas was paraffin-embedded and sectioned into 4μm thick slices by the Translational Pathology Core Laboratory at UCLA.

Immunofluorescence analysis was performed using the Openlab 5.5.0 software on the Leica DM6000 B research microscope. The following antibodies were used: guinea pig anti-insulin (Abcam ab195956, Cambridge, MA, USA, 1:400); mouse anti-glucagon (Sigma-Aldrich G2654, St. Louis, MO, USA, 1:1000); mouse anti-c-Myc (Santa Cruz Biotechnology Inc 9E10 sc-40, Dallas, Texas, USA, 1:100); and mouse anti-HIF1α (Novus Biologicals NB100-105, Centennial, CO, USA, 1:50); The following secondary antibodies were used: F(ab’)2 conjugates with fluorescein isothiocyanate donkey anti-guinea pig immunoglobulin G (IgG) (heavy and light, H + L) (Jackson ImmunoResearch 706-096-148, West Grove, PA, USA, 1:200 for intrinsic factor (IF)) and F(ab’)2 conjugates with Cy3 donkey anti-mouse IgG (H + L) (Jackson ImmunoResearch 711-165-151, West Grove, PA, USA, 1:200 for IF). Vectashield containing 4′,6-diamidino-2-phenylindole (DAPI) (Vector Laboratories H1200, Burlingame, CA, USA) was used to mount the slides. Imaging and data analysis were performed by two observers in a blinded fashion for each section of the experimental mouse genotype.

### Statistical analysis

Data are presented as errors of the means (standard error, SEM) for the number of mice indicated in the figure legends. Mean data were compared between groups by one-way analysis of variance (ANOVA) followed by Tukey’s or Dunnett post-hoc test for multiple comparisons. P-values of less than 0.05 were considered significant.

## Acknowledgments

This manuscript used data acquired from the Human Pancreas Analysis Program (HPAP-RRID:SCR_016202) Database (https://hpap.pmacs.upenn.edu), a Human Islet Research Network (RRID:SCR_014393) consortium (UC4-DK-112217, U01-DK-123594, UC4-DK-112232, and U01-DK-123716). The HPAP study datasets were accessed with appropriate approval through the dbGaP online resource (http://cgems.cancer.gov/data/) (phs002465.v1.p1). This work was supported by funding from Larry Hillblom Foundation (Start-up Grant #2017-D-002-SUP), Hirshberg Foundation Seed Grant HF-2023-024 to ST and Sponsored Research Agreement 2021-0206 between UCLA and Metanoia Bio Inc to ST and KR.

## Authors contribution

ST conceived the paper and designed the experiments. KR, NJ, BS, LS, RP, LY, FM, and JS performed the experiments; FM wrote the module score-based R-script, NJ, LS, RP, LY, and JS performed the bioinformatic analysis, KR, NJ, MP, XY, and ST analyzed the results. ST wrote the manuscript.

## Data and materials availability

All data are available in the main text or the supplementary materials. The custom RScript code with the module score used for this paper will be made available on the GitHub platform.

## Extended Figure Legends

**Figure EV1.**
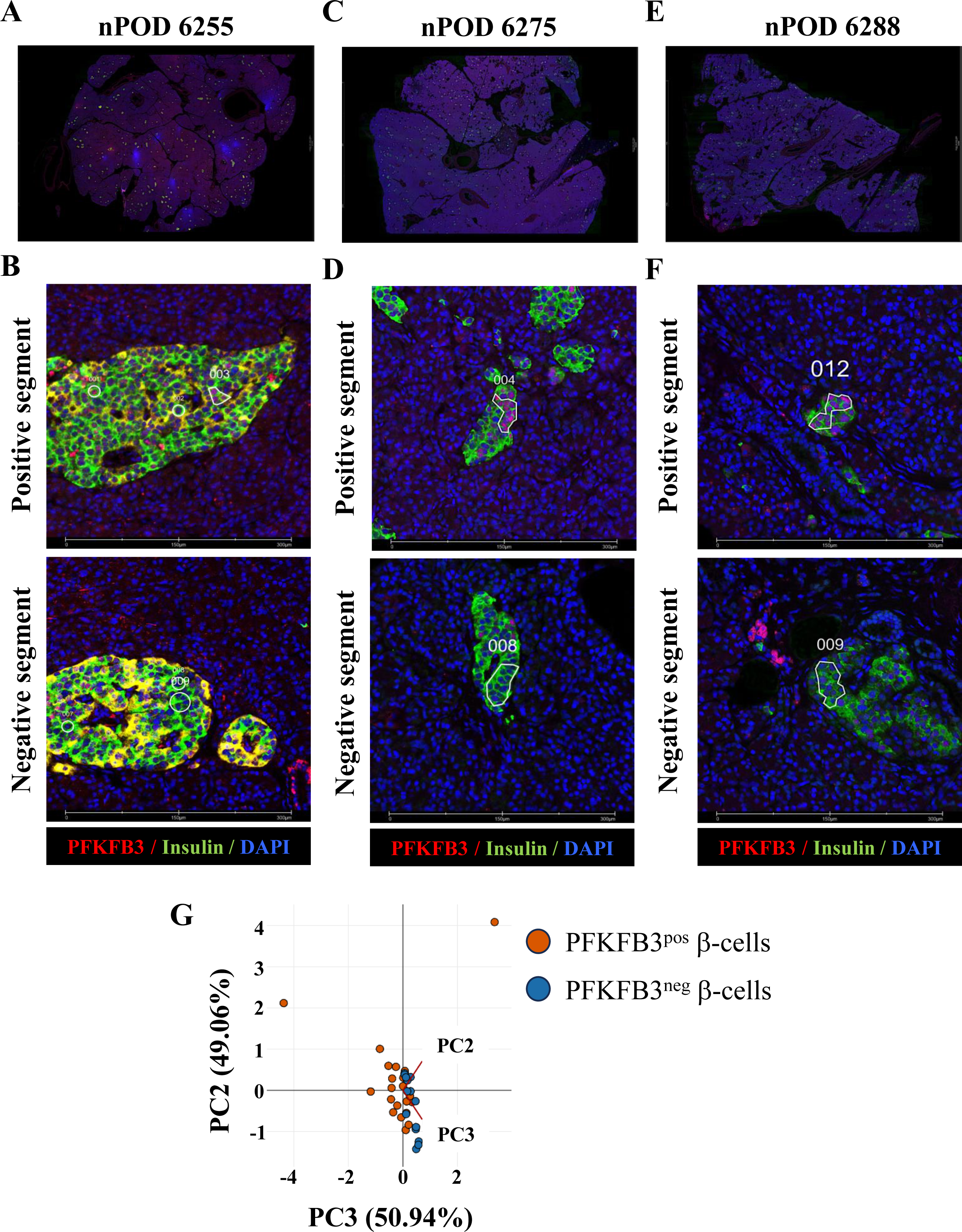
Spatial transcriptomics to transcriptionally profile PFKFB3 positive ;-cells. A-F Whole pancreata sections immunostained with specific antibodies for PFKFB3 (red), Insulin (green) and DAPI (blue) (A, C, E) and PFKFB3 positive (upper panel) and negative (lower panel) segments defining the region of interests (B, D, F) of human nPOD pancreata from T2D donors #6255, #6275, and #6288 respectively. A minimum of 100 nuclei per ROI were segmented and merged for transcriptomic analysis. Scale bar: 150 µm. G The principal component analysis plot for PFKFB3 positive and negative ;-cells

**Figure EV2.**
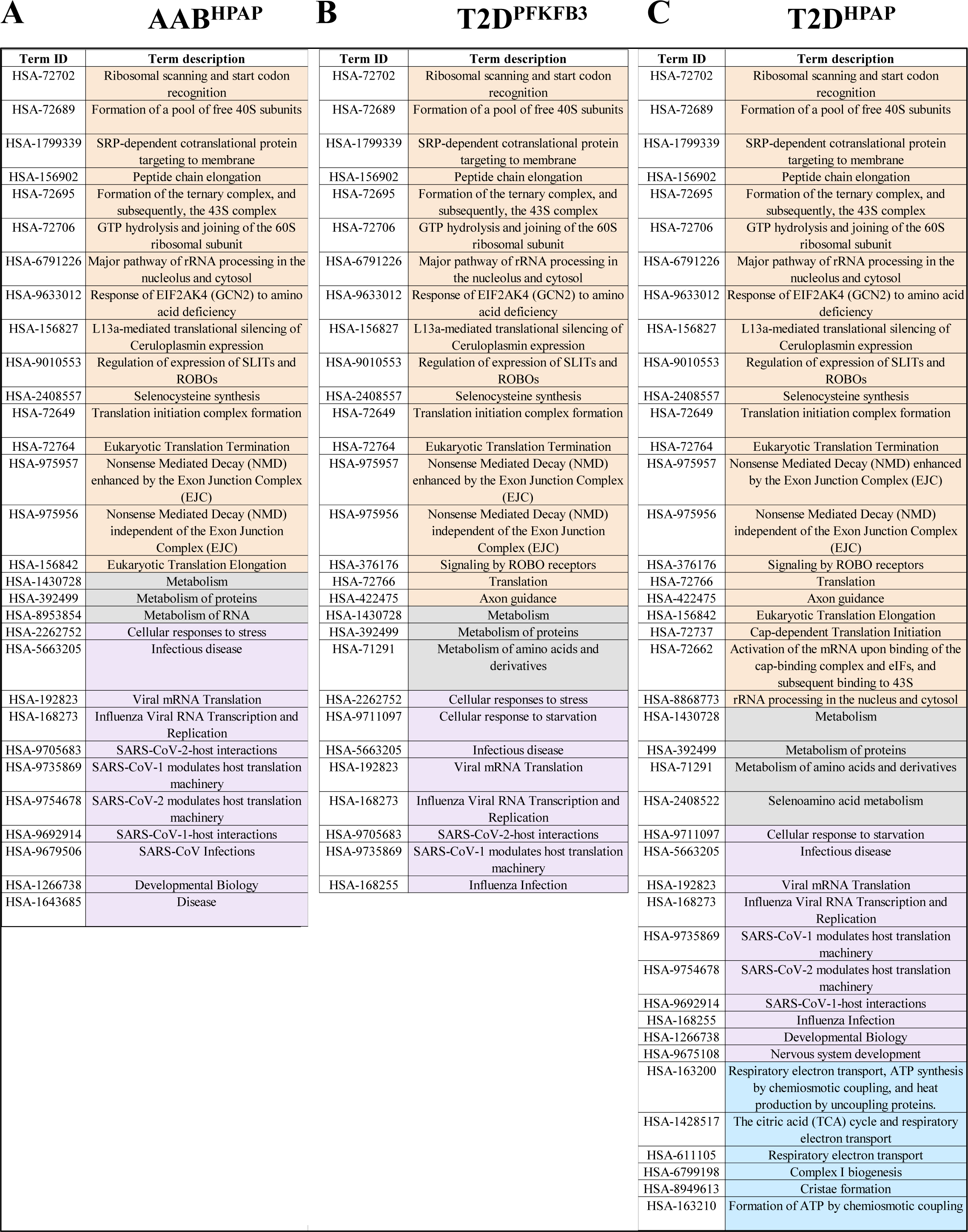
Reactome-based STRING analysis of enriched pathways in different disease states. A-C Tabular presentation of shared enriched pathways between different disease states AAB^HPAP^ (A), T2D^PFKFB3^ (B), and T2D^HPAP^ (C); RiBi pathways are highlighted in orange; mitochondrial respiration pathway is highlighted in blue

**Figure EV3.**
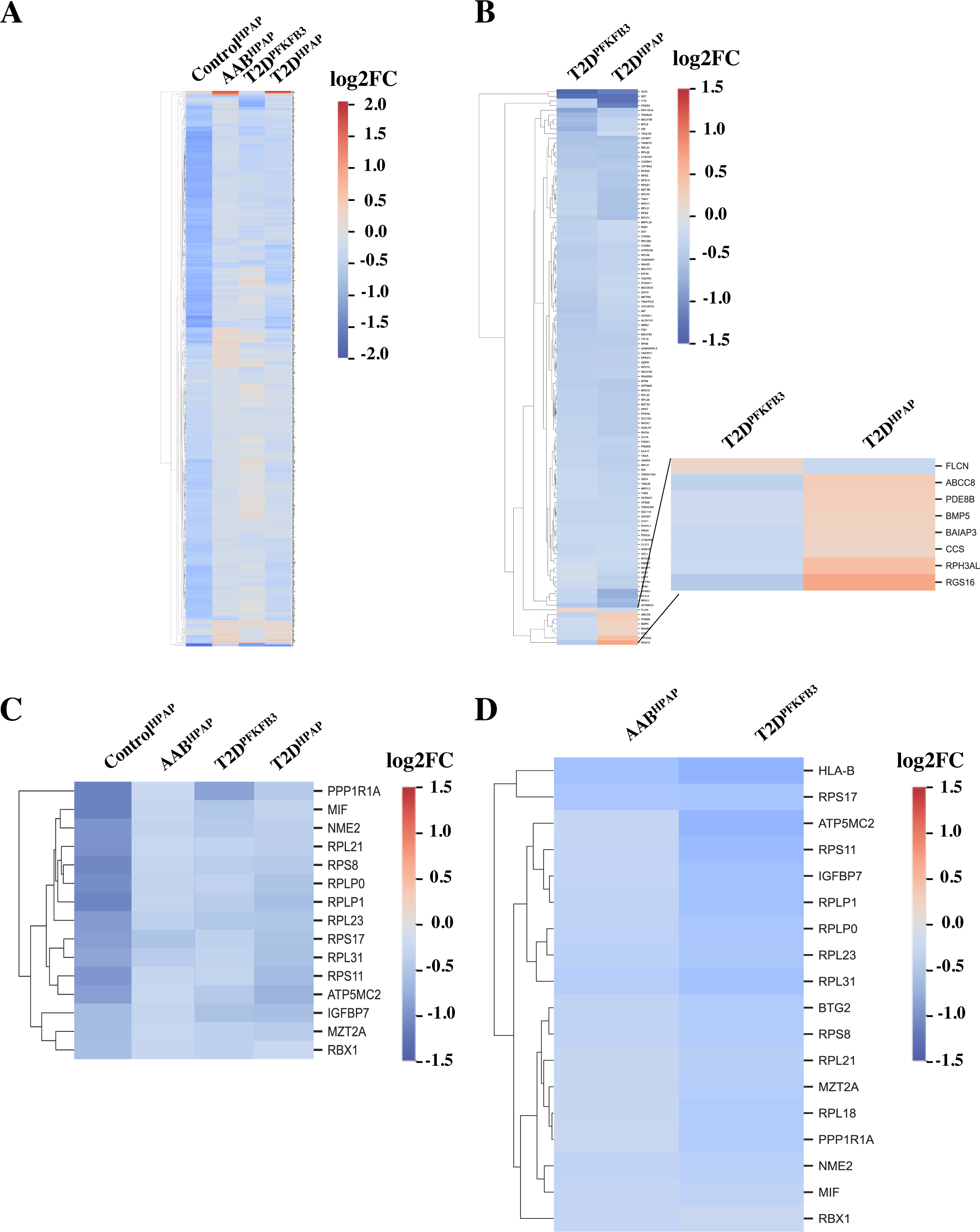
Heatmaps of DEGs from Control^HPAP^, AAB^HPAP^, T2D^PFKFB3^, and T2D^HPAP^. A Heatmap of individual DEGs in Control^HPAP^, AAB^HPAP^, T2D^PFKFB3^, and T2D^HPAP^ B Heatmap of individual DEGs shared by T2D^HPAP^ and T2D^PFKFB3^ with inset showing the inversely correlated genes C Heatmap of individual DEGs (p<0.05) shared by Control^HPAP^, prediabetes-AAB^HPAP^, T2D^HPAP^, and T2D^PFKFB3^ D Heatmap of individual DEGs (p<0.05) shared by AAB^HPAP^ and T2D^PFKFB3^ Scale: log_2_FC

**Figure EV4.**
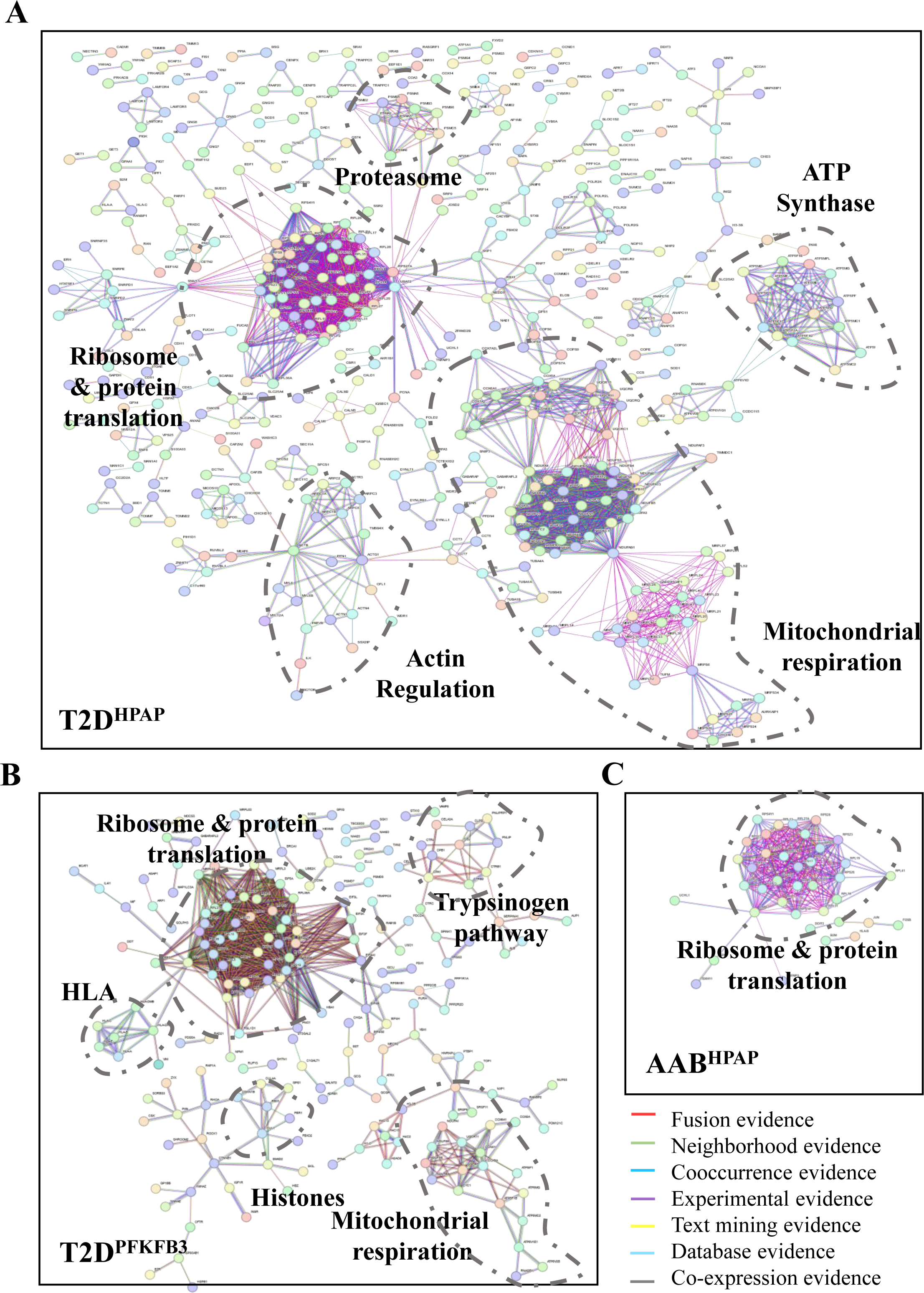
Ribosomes, mitoribosomes, and mitochondrial respiration are at the core of the “loser” signature of PFKFB3-positive ;-cells. A-C Physical evidence (>0.9 confidence level) based protein-protein interaction frameworks were presented with clusters for ribosomal biosynthesis, mitoribosomal genes, and mitochondrial respiration in (A) T2D^HPAP^, (B) T2D^PFKFB3^ and (C) with cluster of ribosomal biosynthesis in AAB^HPAP^. Unique clusters of proteasome, actin regulation, and ATP synthases were revealed in T2D^HPAP^ (A) and trypsinogen pathway, HLA class I & II antigens, and Histones (B) in T2D^PFKFB3^. In the network, the red, green, blue, purple, yellow, light blue, and black lines indicate the presence of fusion evidence, neighborhood evidence, co-occurrence evidence, experimental evidence, text-mining evidence, database evidence, and co-expression evidence, respectively.

**Figure EV5.**
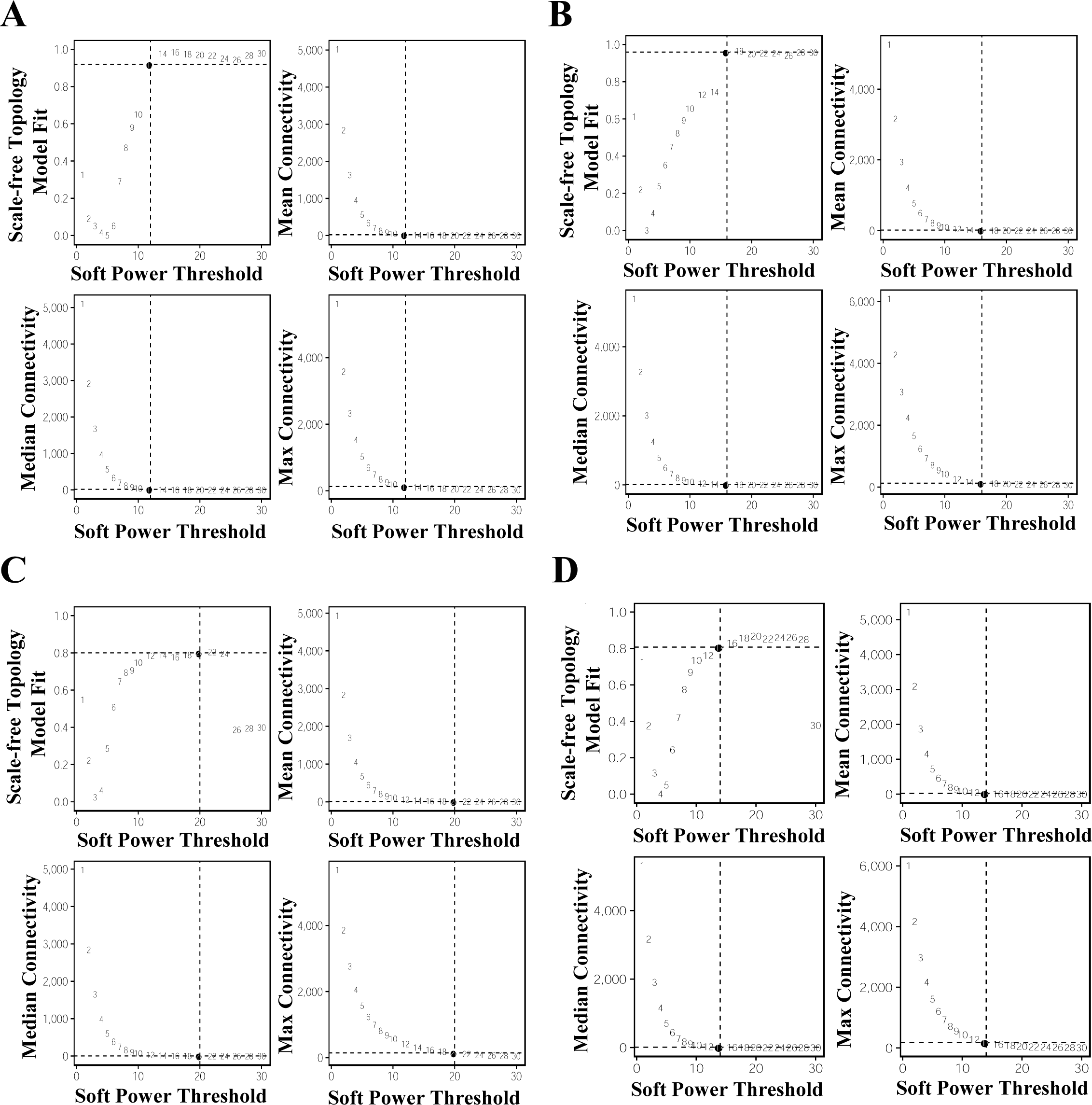
Scale-free topology model fit of hdWGCANA. A-D The soft power threshold was adjusted for each condition to reach scale-free topology in Control^HPAP^ (A), AAB^HPAP^ (B), T1D^HPAP^ (C), and T2D^HPAP^ (D). Mean, median, and maximal connectivity are presented for each condition.

## References

Adachi Y, Sakai T, Terakura S, Shiina T, Suzuki S, Hamana H, Kishi H, Sasazuki T, Arase H, Hanajiri R et al (2022) Downregulation of HLA class II is associated with relapse after allogeneic stem cell transplantation and alters recognition by antigen-specific T cells. Int J Hematol 115: 371–381

Aguayo-Mazzucato C, Andle J, Lee TB, Jr., Midha A, Talemal L, Chipashvili V, Hollister-Lock J, van Deursen J, Weir G, Bonner-Weir S (2019) Acceleration of beta Cell Aging Determines Diabetes and Senolysis Improves Disease Outcomes. Cell Metab 30: 129–142 e124

Arneson D, Bhattacharya A, Shu L, Makinen VP, Yang X (2016) Mergeomics: a web server for identifying pathological pathways, networks, and key regulators via multidimensional data integration. BMC Genomics 17: 722

Baker NE (2020) Emerging mechanisms of cell competition. Nat Rev Genet 21: 683-697

Baumgartner ME, Dinan MP, Langton PF, Kucinski I, Piddini E (2021) Proteotoxic stress is a driver of the loser status and cell competition. Nat Cell Biol 23: 136–146

Blaauw E, van Nieuwenhoven FA, Willemsen P, Delhaas T, Prinzen FW, Snoeckx LH, van Bilsen M, van der Vusse GJ (2010) Stretch-induced hypertrophy of isolated adult rabbit cardiomyocytes. American journal of physiology Heart and circulatory physiology 299: H780–787

Blanco J, Cooper JC, Baker NE (2020) Roles of C/EBP class bZip proteins in the growth and cell competition of Rp (’Minute’) mutants in Drosophila. Elife 9

Bowling S, Lawlor K, Rodriguez TA (2019) Cell competition: the winners and losers of fitness selection. Development 146

Boyd S, Brookfield JL, Critchlow SE, Cumming IA, Curtis NJ, Debreczeni J, Degorce SL, Donald C, Evans NJ, Groombridge S et al (2015) Structure-Based Design of Potent and Selective Inhibitors of the Metabolic Kinase PFKFB3. J Med Chem 58: 3611–3625

Buchan JR, Parker R (2009) Eukaryotic stress granules: the ins and outs of translation. Mol Cell 36: 932–941

Campbell-Thompson M, Wasserfall C, Kaddis J, Albanese-O’Neill A, Staeva T, Nierras C, Moraski J, Rowe P, Gianani R, Eisenbarth G et al (2012) Network for Pancreatic Organ Donors with Diabetes (nPOD): developing a tissue biobank for type 1 diabetes. Diabetes Metab Res Rev 28: 608-617

Cappell SD, Mark KG, Garbett D, Pack LR, Rape M, Meyer T (2018) EMI1 switches from being a substrate to an inhibitor of APC/C(CDH1) to start the cell cycle. Nature 558: 313–317

Carnegie Institution of W (1923) Carnegie Institution of Washington publication. Carnegie Institution of Washington, Washington

Chen W, Wang H, Tao S, Zheng Y, Wu W, Lian F, Jaramillo M, Fang D, Zhang DD (2013) Tumor protein translationally controlled 1 is a p53 target gene that promotes cell survival. Cell Cycle 12: 2321–2328

Chen Y, Chen G (2019) New genetic characteristics of latent autoimmune diabetes in adults (LADA). Ann Transl Med 7: 81

Coelho DS, Moreno E (2019) Emerging links between cell competition and Alzheimer’s disease. J Cell Sci 132

Coelho DS, Schwartz S, Merino MM, Hauert B, Topfel B, Tieche C, Rhiner C, Moreno E (2018) Culling Less Fit Neurons Protects against Amyloid-beta-Induced Brain Damage and Cognitive and Motor Decline. Cell reports 25: 3661-3673 e3663

Cordell HJ (2002) Epistasis: what it means, what it doesn’t mean, and statistical methods to detect it in humans. Hum Mol Genet 11: 2463–2468

de la Cova C, Abril M, Bellosta P, Gallant P, Johnston LA (2004) Drosophila myc regulates organ size by inducing cell competition. Cell 117: 107–116

Ding J, Blencowe M, Nghiem T, Ha SM, Chen YW, Li G, Yang X (2021) Mergeomics 2.0: a web server for multi-omics data integration to elucidate disease networks and predict therapeutics. Nucleic Acids Res 49: W375–W387

Diraison F, Hayward K, Sanders KL, Brozzi F, Lajus S, Hancock J, Francis JE, Ainscow E, Bommer UA, Molnar E et al (2011) Translationally controlled tumour protein (TCTP) is a novel glucose-regulated protein that is important for survival of pancreatic beta cells. Diabetologia 54: 368–379

Diraison F, Ravier MA, Richards SK, Smith RM, Shimano H, Rutter GA (2008) SREBP1 is required for the induction by glucose of pancreatic beta-cell genes involved in glucose sensing. J Lipid Res 49: 814–822

Drake CG, Jaffee E, Pardoll DM (2006) Mechanisms of immune evasion by tumors. Adv Immunol 90: 51–81

Gibson MC, Perrimon N (2005) Extrusion and death of DPP/BMP-compromised epithelial cells in the developing Drosophila wing. Science 307: 1785–1789

Gradeci D, Bove A, Vallardi G, Lowe AR, Banerjee S, Charras G (2021) Cell-scale biophysical determinants of cell competition in epithelia. eLife 10: e61011

Hao Y, Hao S, Andersen-Nissen E, Mauck WM, 3rd, Zheng S, Butler A, Lee MJ, Wilk AJ, Darby C, Zager M et al (2021) Integrated analysis of multimodal single-cell data. Cell 184: 3573-3587 e3529

Huang CJ, Gurlo T, Haataja L, Costes S, Daval M, Ryazantsev S, Wu X, Butler AE, Butler PC (2010) Calcium-activated calpain-2 is a mediator of beta cell dysfunction and apoptosis in type 2 diabetes. J Biol Chem 285: 339–348

Huang da W, Sherman BT, Lempicki RA (2009) Systematic and integrative analysis of large gene lists using DAVID bioinformatics resources. Nat Protoc 4: 44–57

Ji Z, Kiparaki M, Folgado V, Kumar A, Blanco J, Rimesso G, Chuen J, Liu Y, Zheng D, Baker NE (2019) Drosophila RpS12 controls translation, growth, and cell competition through Xrp1. PLoS genetics 15: e1008513

Johnston LA (2009) Competitive interactions between cells: death, growth, and geography. Science 324: 1679–1682

Kaestner KH, Powers AC, Naji A, Consortium H, Atkinson MA (2019) NIH Initiative to Improve Understanding of the Pancreas, Islet, and Autoimmunity in Type 1 Diabetes: The Human Pancreas Analysis Program (HPAP). Diabetes 68: 1394-1402

Kale A, Ji Z, Kiparaki M, Blanco J, Rimesso G, Flibotte S, Baker NE (2018) Ribosomal Protein S12e Has a Distinct Function in Cell Competition. Dev Cell 44: 42–55 e44

Kirschner K, Rattanavirotkul N, Quince MF, Chandra T (2020) Functional heterogeneity in senescence. Biochem Soc Trans 48: 765–773

Konig T, Troder SE, Bakka K, Korwitz A, Richter-Dennerlein R, Lampe PA, Patron M, Muhlmeister M, Guerrero-Castillo S, Brandt U et al (2016) The m-AAA Protease Associated with Neurodegeneration Limits MCU Activity in Mitochondria. Mol Cell 64: 148–162

Kwiatkowska KM, Mavrogonatou E, Papadopoulou A, Sala C, Calzari L, Gentilini D, Bacalini MG, Dall’Olio D, Castellani G, Ravaioli F et al (2023) Heterogeneity of Cellular Senescence: Cell Type-Specific and Senescence Stimulus-Dependent Epigenetic Alterations. Cells 12

Langfelder P, Horvath S (2008) WGCNA: an R package for weighted correlation network analysis. BMC Bioinformatics 9: 559

Lawlor K, Perez-Montero S, Lima A, Rodriguez TA (2020) Transcriptional versus metabolic control of cell fitness during cell competition. Semin Cancer Biol 63: 36–43

Lewis SA, Bakhtiari S, Forstrom J, Bayat A, Bilan F, Le Guyader G, Alkhunaizi E, Vernon H, Padilla-Lopez SR, Kruer MC (2023) AGAP1-associated endolysosomal trafficking abnormalities link gene-environment interactions in a neurodevelopmental disorder. bioRxiv

Leychenko A, Konorev E, Jijiwa M, Matter ML (2011) Stretch-induced hypertrophy activates NFkB-mediated VEGF secretion in adult cardiomyocytes. PloS one 6: e29055

Liaci AM, Steigenberger B, Telles de Souza PC, Tamara S, Grollers-Mulderij M, Ogrissek P, Marrink SJ, Scheltema RA, Forster F (2021) Structure of the human signal peptidase complex reveals the determinants for signal peptide cleavage. Mol Cell 81: 3934–3948 e3911

Lima A, Lubatti G, Burgstaller J, Hu D, Green AP, Di Gregorio A, Zawadzki T, Pernaute B, Mahammadov E, Perez-Montero S et al (2021) Cell competition acts as a purifying selection to eliminate cells with mitochondrial defects during early mouse development. Nat Metab 3: 1091–1108

Lucchesi JC (1968) Synthetic lethality and semi-lethality among functionally related mutants of Drosophila melanfgaster. Genetics 59: 37–44

MacLean RC, Perron GG, Gardner A (2010) Diminishing returns from beneficial mutations and pervasive epistasis shape the fitness landscape for rifampicin resistance in Pseudomonas aeruginosa. Genetics 186: 1345–1354

Melber A, Haynes CM (2018) UPR(mt) regulation and output: a stress response mediated by mitochondrial-nuclear communication. Cell Res 28: 281–295

Merino MM, Levayer R, Moreno E (2016) Survival of the Fittest: Essential Roles of Cell Competition in Development, Aging, and Cancer. Trends in cell biology 26: 776–788

Min J, Ma F, Seyran B, Pellegrini M, Greeff O, Moncada S, Tudzarova S (2022) beta-cell-specific deletion of PFKFB3 restores cell fitness competition and physiological replication under diabetogenic stress. Commun Biol 5: 248

Moiseeva V, Cisneros A, Sica V, Deryagin O, Lai Y, Jung S, Andres E, An J, Segales J, Ortet L et al (2022) Senescence atlas reveals an aged-like inflamed niche that blunts muscle regeneration. Nature

Montemurro C, Nomoto H, Pei L, Parekh VS, Vongbunyong KE, Vadrevu S, Gurlo T, Butler AE, Subramaniam R, Ritou E et al (2019) IAPP toxicity activates HIF1alpha/PFKFB3 signaling delaying beta-cell loss at the expense of beta-cell function. Nat Commun 10: 2679

Morabito S, Reese F, Rahimzadeh N, Miyoshi E, Swarup V (2023) hdWGCNA identifies co-expression networks in high-dimensional transcriptomics data. Cell reports methods 3: 100498

Moreno E, Basler K, Morata G (2002) Cells compete for decapentaplegic survival factor to prevent apoptosis in Drosophila wing development. Nature 416: 755–759

Munch C (2018) The different axes of the mammalian mitochondrial unfolded protein response. BMC Biol 16: 81

Nomoto H, Pei L, Montemurro C, Rosenberger M, Furterer A, Coppola G, Nadel B, Pellegrini M, Gurlo T, Butler PC et al (2020) Activation of the HIF1alpha/PFKFB3 stress response pathway in beta cells in type 1 diabetes. Diabetologia 63: 149–161

O’Neil NJ, Bailey ML, Hieter P (2017) Synthetic lethality and cancer. Nature Reviews Genetics 18: 613–623

Opalinska M, Janska H (2018) AAA Proteases: Guardians of Mitochondrial Function and Homeostasis. Cells 7

Ortiz-Barahona A, Villar D, Pescador N, Amigo J, del Peso L (2010) Genome-wide identification of hypoxia-inducible factor binding sites and target genes by a probabilistic model integrating transcription-profiling data and in silico binding site prediction. Nucleic Acids Res 38: 2332–2345

Perfeito L, Sousa A, Bataillon T, Gordo I (2014) Rates of fitness decline and rebound suggest pervasive epistasis. Evolution 68: 150–162

Ramirez Reyes JMJ, Cuesta R, Pause A (2021) Folliculin: A Regulator of Transcription Through AMPK and mTOR Signaling Pathways. Front Cell Dev Biol 9: 667311

Richter JD, Coller J (2015) Pausing on Polyribosomes: Make Way for Elongation in Translational Control. Cell 163: 292–300

Rosario FJ, Powell TL, Gupta MB, Cox L, Jansson T (2020) mTORC1 Transcriptional Regulation of Ribosome Subunits, Protein Synthesis, and Molecular Transport in Primary Human Trophoblast Cells. Front Cell Dev Biol 8: 583801

Sasai N, Agata N, Inoue-Miyazu M, Kawakami K, Kobayashi K, Sokabe M, Hayakawa K (2010) Involvement of PI3K/Akt/TOR pathway in stretch-induced hypertrophy of myotubes. Muscle & nerve 41: 100–106

Schoustra S, Hwang S, Krug J, de Visser JA (2016) Diminishing-returns epistasis among random beneficial mutations in a multicellular fungus. Proc Biol Sci 283

Shannon P, Markiel A, Ozier O, Baliga NS, Wang JT, Ramage D, Amin N, Schwikowski B, Ideker T (2003) Cytoscape: a software environment for integrated models of biomolecular interaction networks. Genome Res 13: 2498–2504

Sherman BT, Hao M, Qiu J, Jiao X, Baseler MW, Lane HC, Imamichi T, Chang W (2022) DAVID: a web server for functional enrichment analysis and functional annotation of gene lists (2021 update). Nucleic Acids Res 50: W216-W221

Shetty S, Hofstetter J, Battaglioni S, Ritz D, Hall MN (2023) TORC1 phosphorylates and inhibits the ribosome preservation factor Stm1 to activate dormant ribosomes. EMBO J 42: e112344

Shi FD, Ljunggren HG, La Cava A, Van Kaer L (2011) Organ-specific features of natural killer cells. Nat Rev Immunol 11: 658–671

Shiravand Y, Khodadadi F, Kashani SMA, Hosseini-Fard SR, Hosseini S, Sadeghirad H, Ladwa R, O’Byrne K, Kulasinghe A (2022) Immune Checkpoint Inhibitors in Cancer Therapy. Curr Oncol 29: 3044–3060

Szklarczyk D, Gable AL, Nastou KC, Lyon D, Kirsch R, Pyysalo S, Doncheva NT, Legeay M, Fang T, Bork P et al (2021a) Correction to ‘The STRING database in 2021: customizable protein-protein networks, and functional characterization of user-uploaded gene/measurement sets’. Nucleic Acids Res 49: 10800

Szklarczyk D, Gable AL, Nastou KC, Lyon D, Kirsch R, Pyysalo S, Doncheva NT, Legeay M, Fang T, Bork P et al (2021b) The STRING database in 2021: customizable protein-protein networks, and functional characterization of user-uploaded gene/measurement sets. Nucleic acids research 49: D605–d612

Tamori Y, Deng WM (2014) Compensatory cellular hypertrophy: the other strategy for tissue homeostasis. Trends in cell biology 24: 230–237

Thompson PJ, Shah A, Ntranos V, Van Gool F, Atkinson M, Bhushan A (2019) Targeted Elimination of Senescent Beta Cells Prevents Type 1 Diabetes. Cell Metab 29: 1045–1060 e1010

Traxler L, Herdy JR, Stefanoni D, Eichhorner S, Pelucchi S, Szucs A, Santagostino A, Kim Y, Agarwal RK, Schlachetzki JCM et al (2022) Warburg-like metabolic transformation underlies neuronal degeneration in sporadic Alzheimer’s disease. Cell Metab 34: 1248–1263 e1246

Tudzarova S, Colombo SL, Stoeber K, Carcamo S, Williams GH, Moncada S (2011) Two ubiquitin ligases, APC/C-Cdh1 and SKP1-CUL1-F (SCF)-beta-TrCP, sequentially regulate glycolysis during the cell cycle. Proc Natl Acad Sci U S A 108: 5278-5283

Tutaj H, Tomala K, Korona R, 2023. Epistasis supports viability under extensive gene-dose insufficiency following chromosome loss. Cold Spring Harbor Laboratory.

Valvona CJ, Fillmore HL, Nunn PB, Pilkington GJ (2016) The Regulation and Function of Lactate Dehydrogenase A: Therapeutic Potential in Brain Tumor. Brain Pathol 26: 3–17

Vieira R, Mariani JN, Huynh NPT, Stephensen HJT, Solly R, Tate A, Schanz S, Cotrupi N, Mousaei M, Sporring J et al (2023) Young glial progenitor cells competitively replace aged and diseased human glia in the adult chimeric mouse brain. Nature Biotechnology

Wallace C, Cutler AJ, Pontikos N, Pekalski ML, Burren OS, Cooper JD, Garcia AR, Ferreira RC, Guo H, Walker NM et al (2015) Dissection of a Complex Disease Susceptibility Region Using a Bayesian Stochastic Search Approach to Fine Mapping. PLoS Genet 11: e1005272

Wheeler JR, Matheny T, Jain S, Abrisch R, Parker R (2016) Distinct stages in stress granule assembly and disassembly. Elife 5

Wigger L, Barovic M, Brunner AD, Marzetta F, Schoniger E, Mehl F, Kipke N, Friedland D, Burdet F, Kessler C et al (2021) Multi-omics profiling of living human pancreatic islet donors reveals heterogeneous beta cell trajectories towards type 2 diabetes. Nat Metab 3: 1017–1031

