## Supplementary Figures S1 to S7 for "Dysfunctional β-cell longevity in diabetes relies on energy conservation and positive epistasis"

**Appendix -1**

**Table of contents**

1. Legends for supplementary figures
2. Supplementary figures S1 to S7

#### 1. Legends for supplementary figures

**Figure S1 – Strength of protein-protein interactions**

Dot plot showing the strength ofSupplementary protein-protein interactions in Reactome enriched pathways shared between $AAB^{HPAP}$, $T2D^{HPAP}$ and $T2D^{PFKFB3}$.

**Figure S2 - Ribosomal biosynthesis (RiBi) and protein translation are shared pathways between β-cells with "loser signature" in $AAB^{HPAP}$, $T2D^{HPAP}$ and PFKFB3-positive β-cells in T2D ($T2D^{PFKFB3}$)**

A-C Pair-wise overlap of enriched pathways between $T2D^{HPAP}$ and $T2D^{PFKFB3}$ (A), $T2D^{HPAP}$ and $AAB^{HPAP}$ (B) and $T2D^{PFKFB3}$ and $AAB^{HPAP}$ (C). In the presented networks, thicker edges between enriched pathways indicate a higher significance and lower p-value, while colors such as red depict a positive association and grey a negative association.

**Figure S3 – Feature plots of different metacell modules for "loser" β-cells generated by hdWGCNA**

A-D Feature plots of different metacell modules for "loser" β-cells $Control^{HPAP}$ (A), $AAB^{HPAP}$ (B), $T1D^{HPAP}$ (C), and $T2D^{HPAP}$ (D); the scale represents Z-score.

**Figure S4 – Correlation between metacell modules in hdWGCNA**

A-D Correlation between metacell modules for "loser" β-cells in $Control^{HPAP}$ (A), $AAB^{HPAP}$ (B), $T1D^{HPAP}$ (C), and $T2D^{HPAP}$ (D); the scale represents Z-score.

**Figure S5 - Individual and merged immunofluorescence images for bihormonal cells in Fig 5E**

Representative individual and merged immunofluorescence images of islets from wild-type + high fat diet (WT), DS + vehicle, $PFKFB3^{\beta KO}$ DS, and DS + iPFKFB3 mice, immunostained for glucagon (red), insulin (green) and nuclei (blue) showing eradication of glucagon and insulin positive injured bihormonal cells in $PFKFB3^{\beta KO}$ DS and DS + iPFKFB3 mice.

**Figure S6 - Individual and merged immunofluorescence images for cytoplasmic c-Myc positive cells in Fig 5E**

Representative individual and merged immunofluorescence images of islets from wild-type + high fat diet (WT), DS + vehicle, $PFKFB3^{\beta KO}$ DS, and DS + iPFKFB3 mice, immunostained for cytoplasmic c-Myc (red), insulin (green) and nuclei (blue) showing eradication of c-Myc positive injured β-cells in $PFKFB3^{\beta KO}$ DS and DS + iPFKFB3 mice

**Figure S7 - Individual and merged immunofluorescence images for HIF1α positive cells in Fig 5E**

Representative individual and merged immunofluorescence images of islets from wild-type + high fat diet (WT), DS + vehicle, PFKFB3$^{\beta KO}$ DS, and DS + iPFKFB3 mice, immunostained for HIF1α (red), insulin (green) and nuclei (blue) showing eradication of HIF1α - positive injured β-cells in PFKFB3$^{\beta KO}$ DS and DS + iPFKFB3 mice

### Figure S1

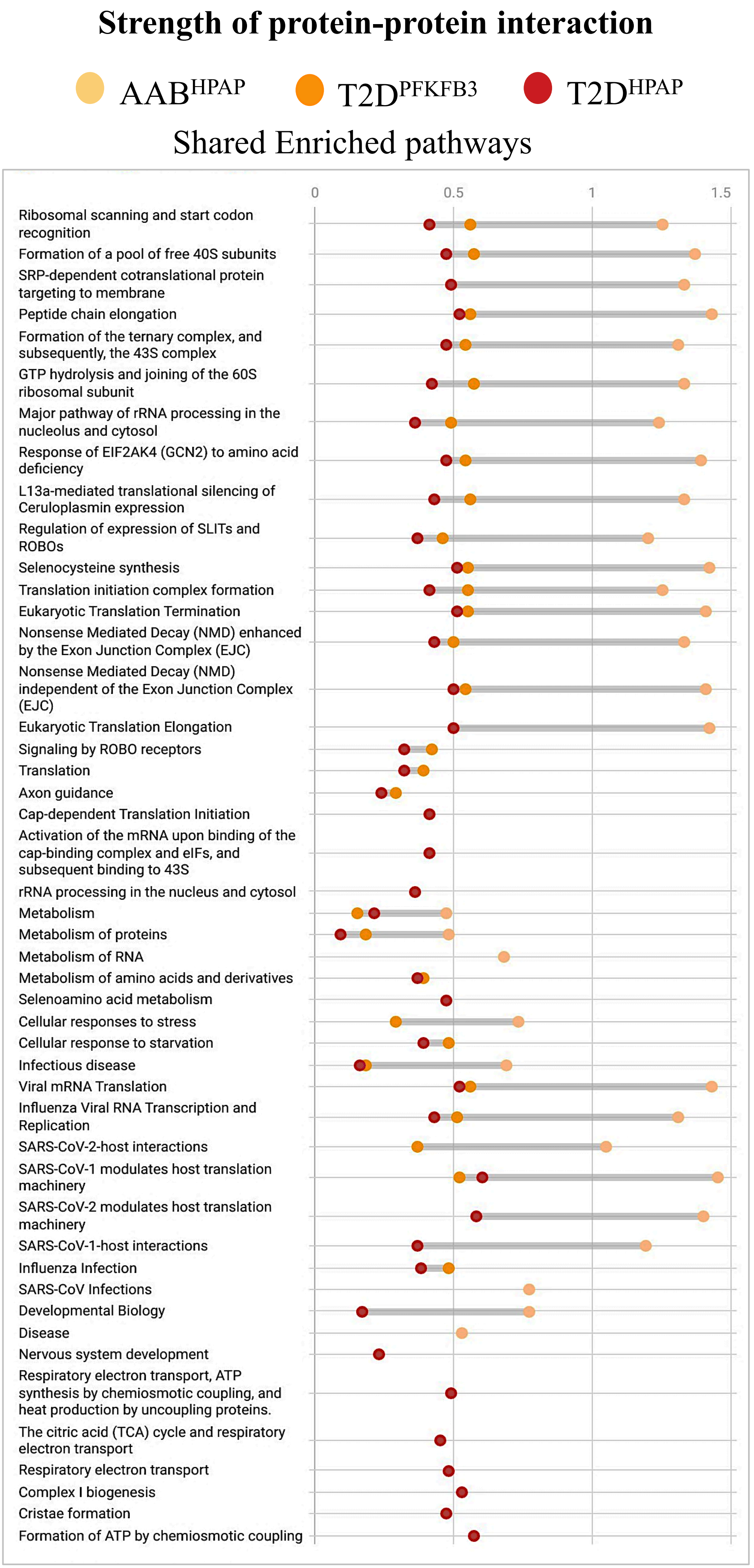
Strength of protein-protein interaction
AAB$^{HPAP}$
T2D$^{PFKFB3}$
T2D$^{HPAP}$
Shared Enriched pathways
0
0.5
1
1.5
Ribosomal scanning and start codon recognition
Formation of a pool of free 40S subunits
SRP-dependent cotranslational protein targeting to membrane
Peptide chain elongation
Formation of the ternary complex, and subsequently, the 43S complex
GTP hydrolysis and joining of the 60S ribosomal subunit
Major pathway of rRNA processing in the nucleolus and cytosol
Response of EIF2AK4 (GCN2) to amino acid deficiency
L13a-mediated translational silencing of Ceruloplasmin expression
Regulation of expression of SLITs and ROBOs
Selenocysteine synthesis
Translation initiation complex formation
Eukaryotic Translation Termination
Nonsense Mediated Decay (NMD) enhanced by the Exon Junction Complex (EJC)
Nonsense Mediated Decay (NMD) independent of the Exon Junction Complex (EJC)
Eukaryotic Translation Elongation
Signaling by ROBO receptors
Translation
Axon guidance
Cap-dependent Translation Initiation
Activation of the mRNA upon binding of the cap-binding complex and eIFs, and subsequent binding to 43S
rRNA processing in the nucleus and cytosol
Metabolism
Metabolism of proteins
Metabolism of RNA
Metabolism of amino acids and derivatives
Selenoamino acid metabolism
Cellular responses to stress
Cellular response to starvation
Infectious disease
Viral mRNA Translation
Influenza Viral RNA Transcription and Replication
SARS-CoV-2-host interactions
SARS-CoV-1 modulates host translation machinery
SARS-CoV-2 modulates host translation machinery
SARS-CoV-1-host interactions
Influenza Infection
SARS-CoV Infections
Developmental Biology
Disease
Nervous system development
Respiratory electron transport, ATP synthesis by chemiosmotic coupling, and heat production by uncoupling proteins.
The citric acid (TCA) cycle and respiratory electron transport
Respiratory electron transport
Complex I biogenesis
Cristae formation
Formation of ATP by chemiosmotic coupling

### Figure S2

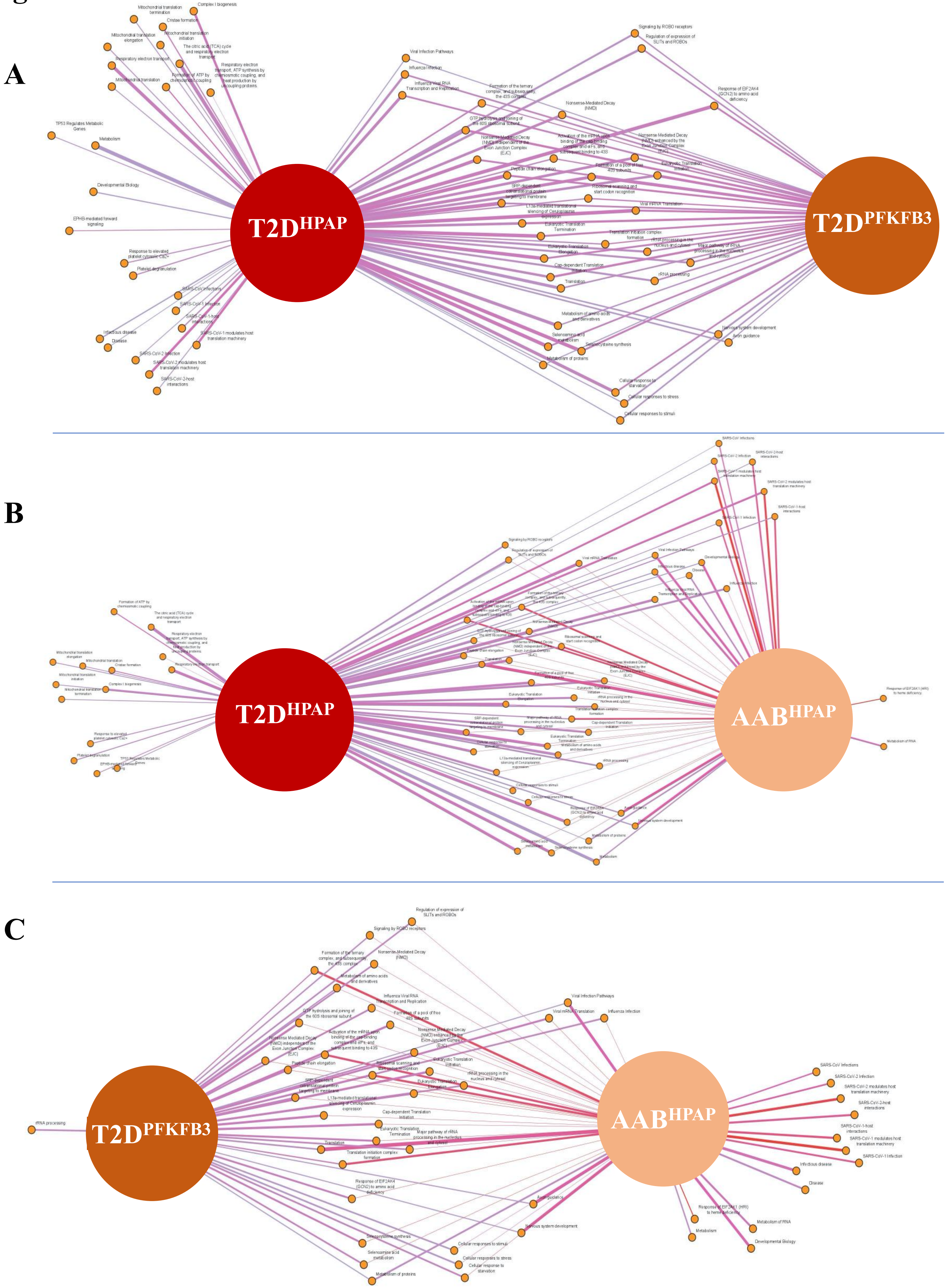

### Figure S3

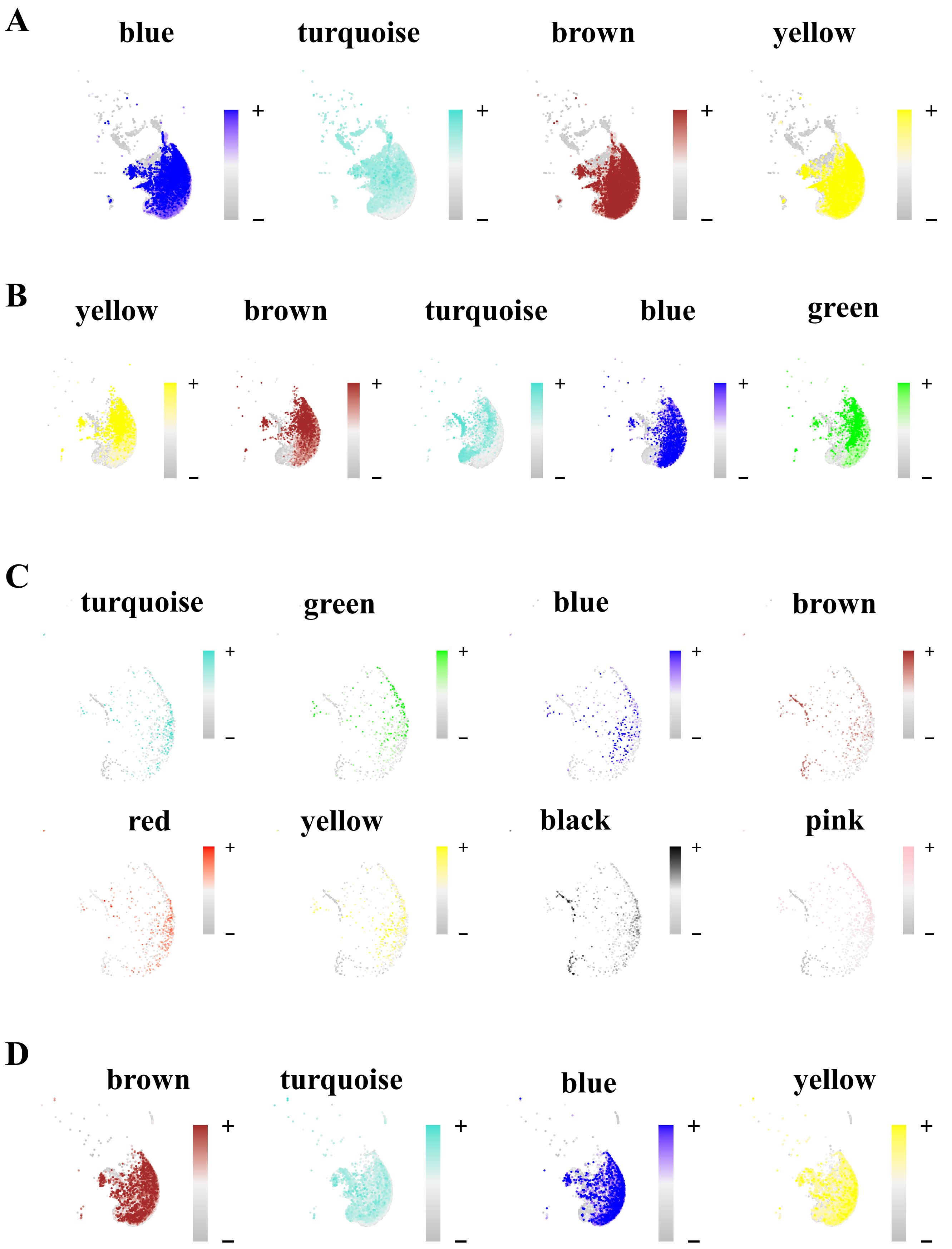

A
blue
turquoise
brown
yellow
B
yellow
brown
turquoise
blue
green
C
turquoise
green
blue
brown
red
yellow
black
pink
D
brown
turquoise
blue
yellow

### Figure S4

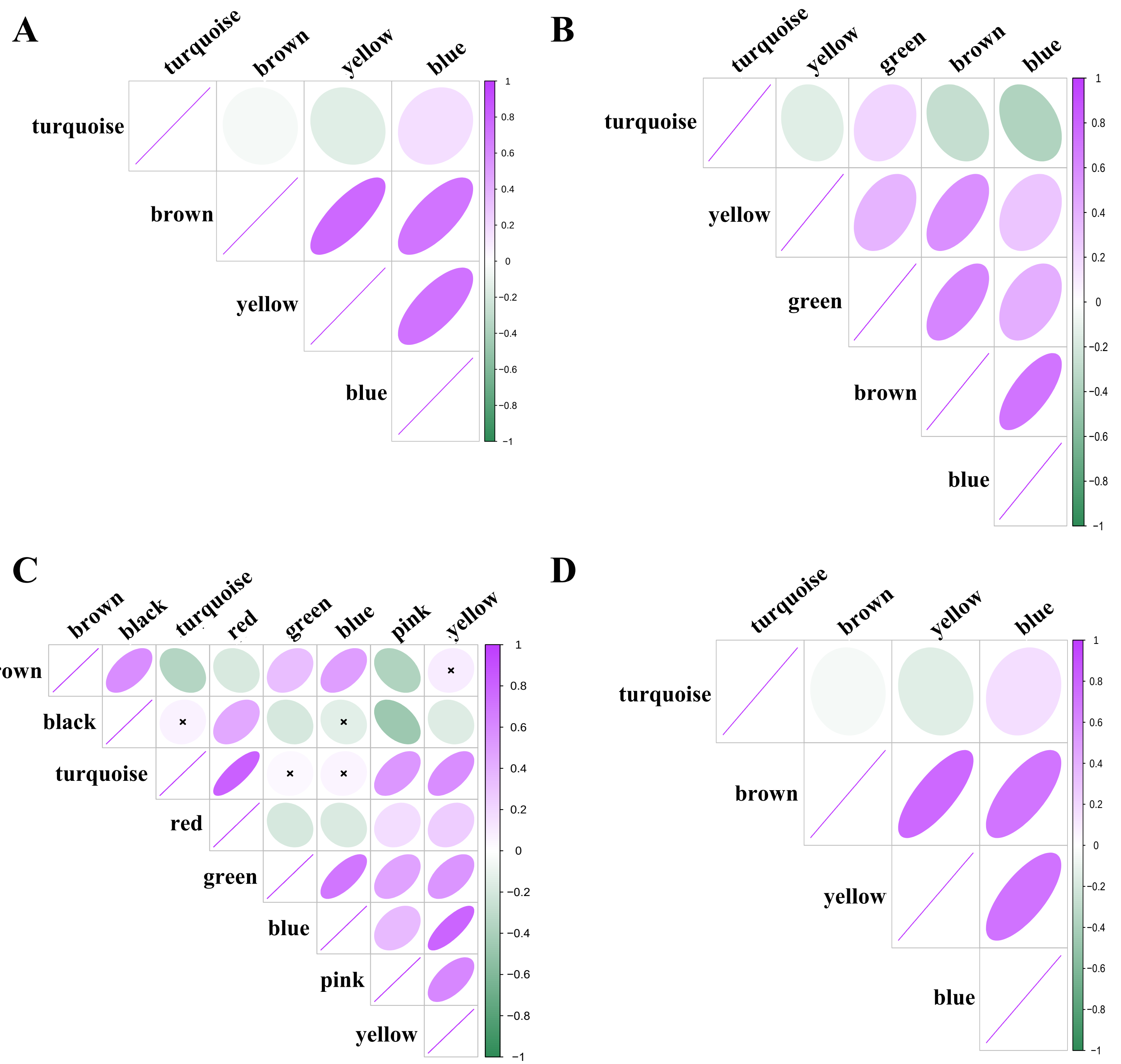
A
turquoise
brown
yellow
blue
1
0.8
0.6
0.4
0.2
0
−0.2
−0.4
−0.6
−0.8
−1
B
turquoise
yellow
green
brown
blue
C
brown
black
turquoise
red
green
blue
pink
yellow
D
turquoise
brown
yellow
blue

### Figure S5

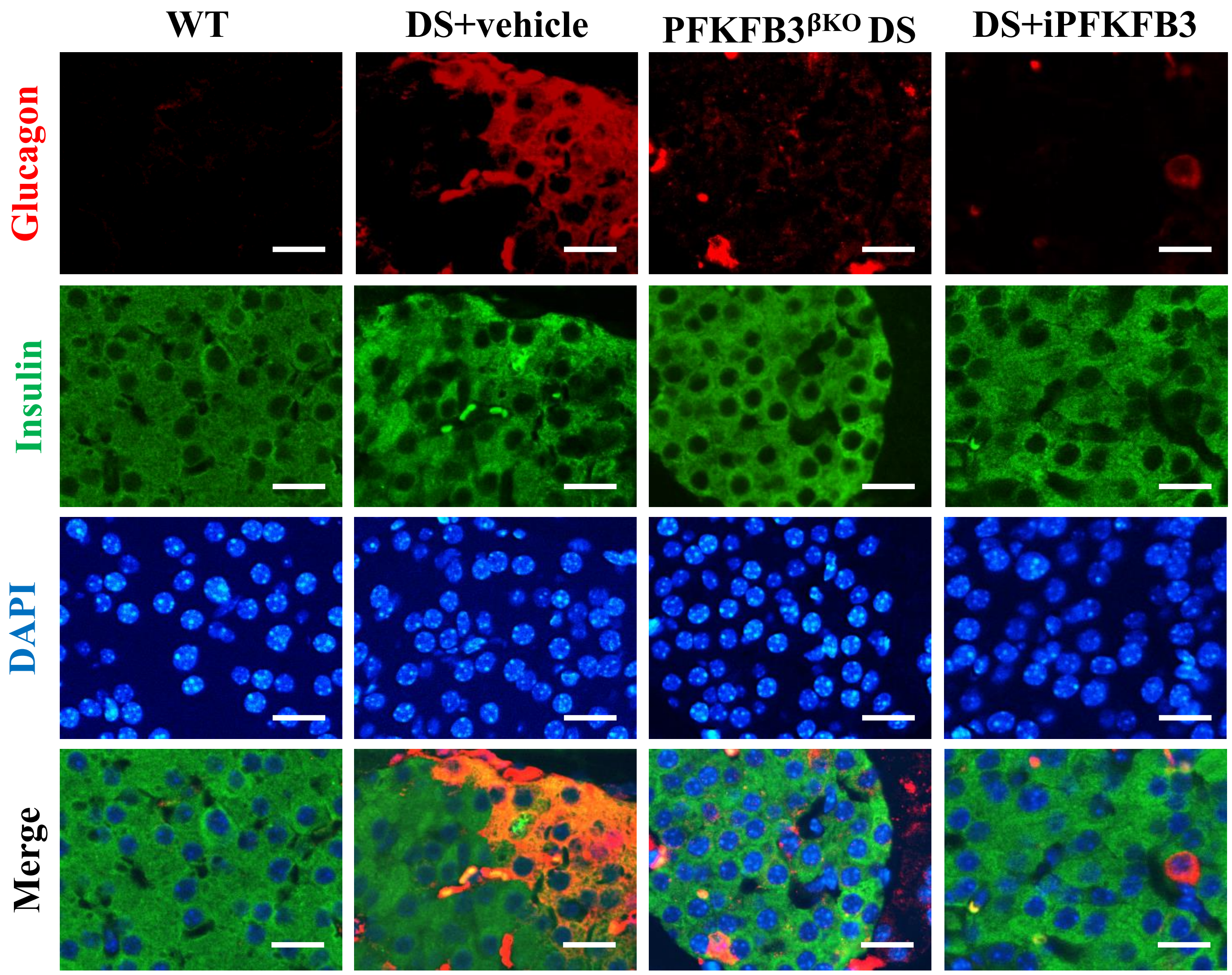
WT
DS+vehicle
PFKFB3βKO DS
DS+iPFKFB3
Glucagon
Insulin
DAPI
Merge

### Figure S6

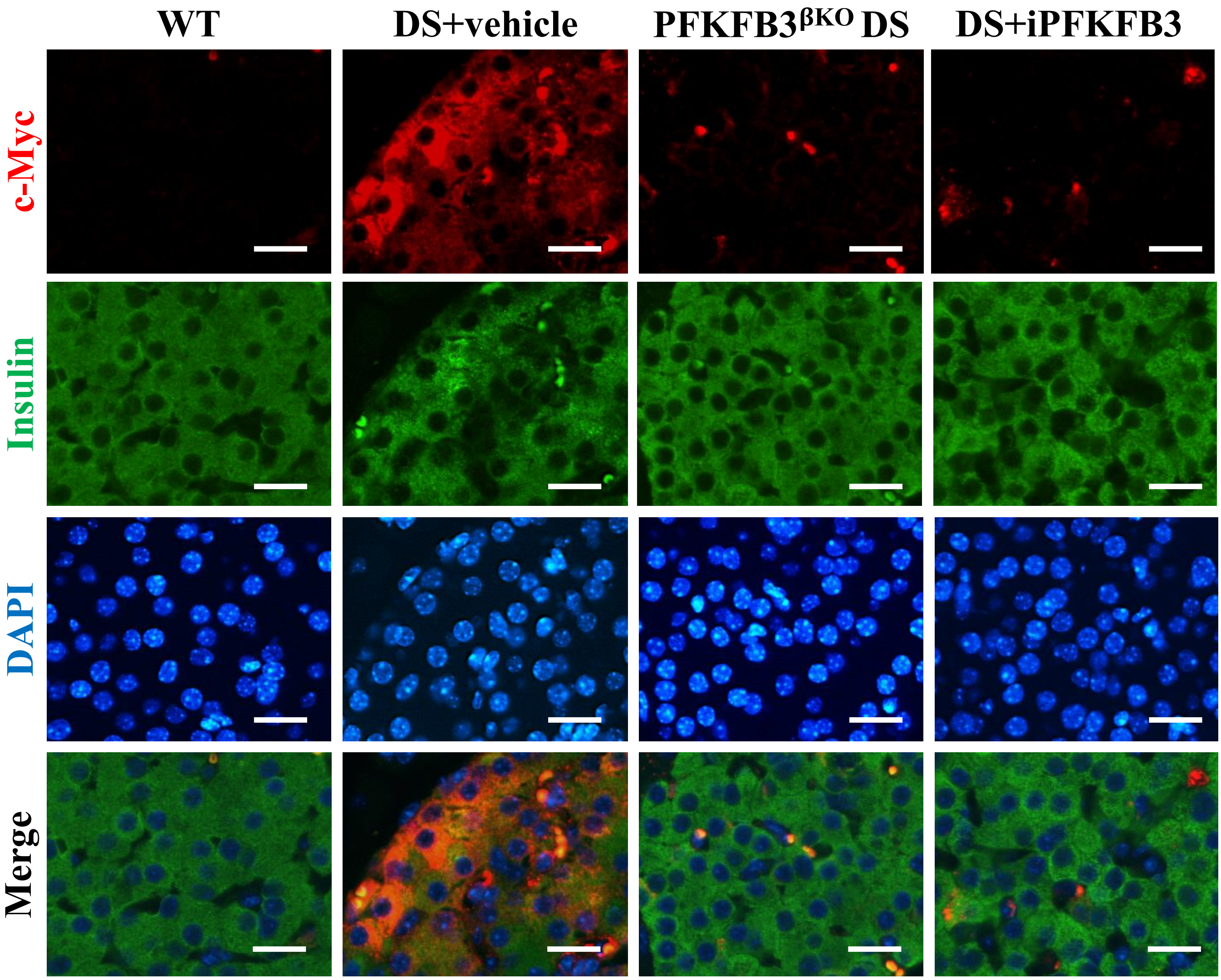
WT
DS+vehicle
PFKFB3βKO DS
DS+iPFKFB3
c-Myc
Insulin
DAPI
Merge

### Figure S7

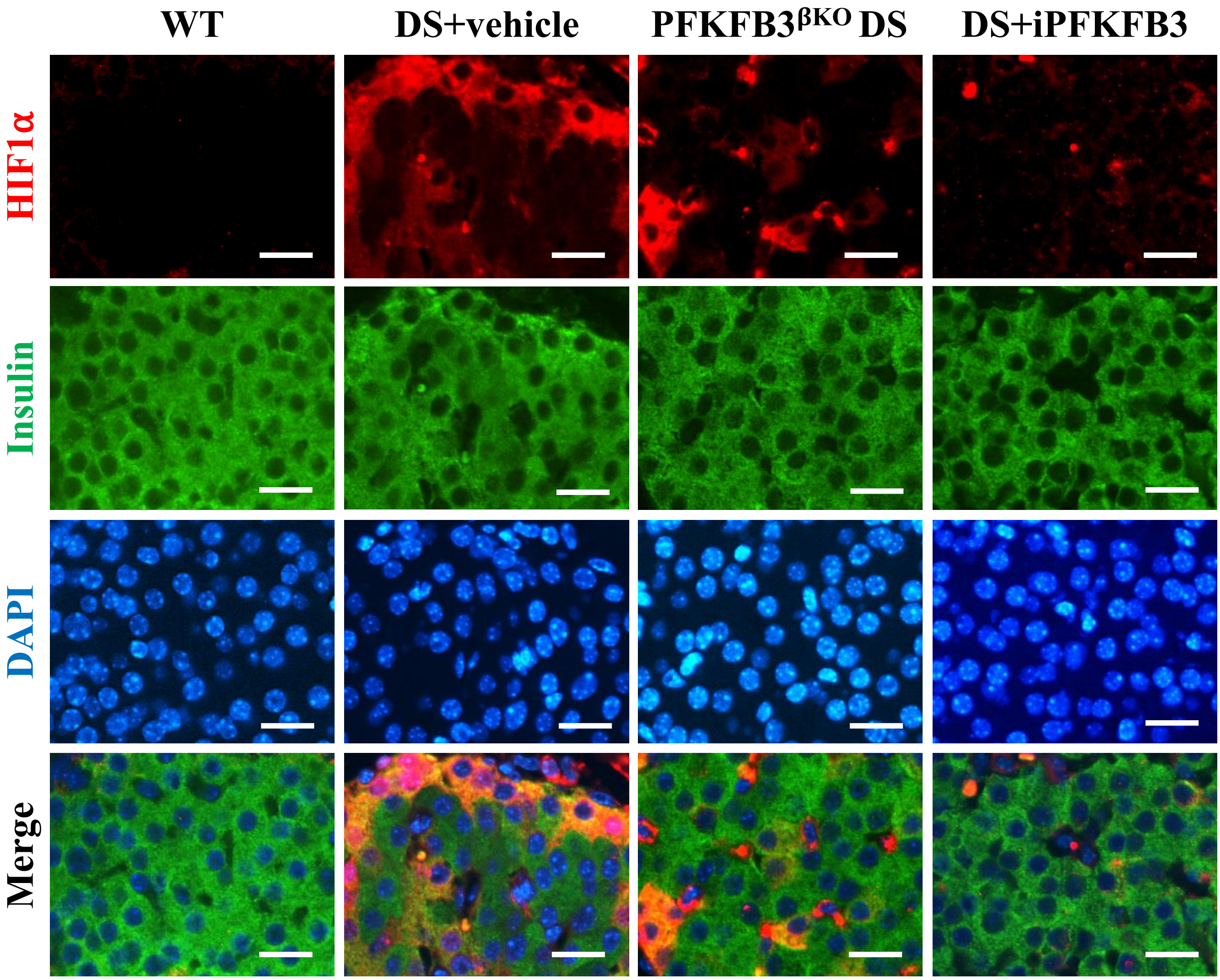
WT
DS+vehicle
PFKFB3βKO DS
DS+iPFKFB3
HIF1α
Insulin
DAPI
Merge
